# Comprehensive analysis of meiosis-derived cDNA libraries reveals gene isoforms and mitochondrial proteins important for competitive fitness

**DOI:** 10.1101/2021.05.10.443361

**Authors:** Tina L. Sing, Katie Conlon, Stephanie H. Lu, Nicole Madrazo, Juliet C. Barker, Ina Hollerer, Gloria A. Brar, Peter H. Sudmant, Elçin Ünal

**Affiliations:** Department of Molecular and Cell Biology, University of California, Berkeley, CA, USA; Department of Integrative Biology, University of California, Berkeley, CA, USA

## Abstract

Gametogenesis is a highly regulated and dynamic developmental program where a diploid progenitor cell differentiates into haploid gametes, the precursors for sexual reproduction. During meiosis, several pathways converge to initiate ploidy reduction and organelle remodelling to render gametes competent for zygote formation and subsequent organismal development. Additionally, meiosis inherently rejuvenates the newly formed gametes resulting in lifespan resetting. Here, we construct five stage-specific, inducible meiotic cDNA libraries that represent over 84% of the yeast genome. We employ computational strategies to detect stage-specific meiotic transcript isoforms in each library and develop a robust screening pipeline to test the effect of each cDNA on competitive fitness. Our multi-day proof-of-principle time course reveals gene isoforms that are important for competitive fitness as well as mitochondrial proteins that cause dose-dependent disruption of respiration. Together, these novel meiotic cDNA libraries provide an important resource for systematically studying meiotic genes and gene isoforms in future studies.

**HIGHLIGHTS:**

- Construction of five stage-specific, inducible meiotic cDNA libraries in budding yeast that collectively represent 5563 genes, which is over 84% of the genome
- Analysis of the cDNA libraries reveal the presence of meiosis-specific transcript isoforms that are largely uncharacterized
- Development of a robust gain-of-function screening pipeline identifies previously characterized genes and novel gene isoforms important for competitive fitness
- Multi-day proof-of-principle screen reveals mitochondrial proteins that cause dosage-specific respiration defects

## INTRODUCTION

For a species to persist, organisms must fulfill the biological imperative to reproduce and create new offspring. In eukaryotes, this is commonly accomplished through sexual reproduction, the biological process by which the genetic information from two living organisms is combined to generate progeny that are genetically distinct from either parent. This form of reproduction relies on the formation of haploid gametes, known as “egg” and “sperm” in metazoans, through a highly regulated and dynamic developmental program called gametogenesis. During this process, a diploid progenitor cell must: (i) undergo meiotic divisions that halve the genome, (ii) accurately partition and synthesize organelles; and (iii) exclude or eliminate age-associated biomarkers from the developing gametes (Bolcun-Filas and Handel, 2018; Goodman et al., 2020; King and Ünal, 2020; Marston and Amon, 2004). Together, these pathways converge to produce gametes that are competent for zygote formation and subsequent organismal development.

Because gametogenesis is highly conserved among eukaryotes, significant progress in understanding gamete formation has been made by studying the analogous spore formation in the single-celled eukaryote, *Saccharomyces cerevisiae* (Neiman, 2011). In the wild, budding yeast gametogenesis, also referred to as sporulation, is naturally induced by nutrient deprivation to promote prolonged survival of gametes until nutrients become available again (Freese et al., 1982). As yeast enter a starvation state, the decision to transition from the mitotic to meiotic fate is initiated by a key transcription factor Ime1 (Kassir et al., 1988). This leads to entrance into early meiosis where DNA replication, double strand break (DSB) formation, pairing of homologous chromosomes, synaptonemal complex assembly, and the initiation of recombination occurs. Importantly, Ime1 also induces the expression of a second key transcription factor, Ndt80, which is related to the p53 family (Xu et al., 1995). Repair of DSBs leads to Ndt80 activation, which triggers two rounds of meiotic divisions, termed “Meiosis I” and “Meiosis II”, culminating in four haploid nuclei lobes that are surrounded by developing gamete plasma membrane that is made *de novo* from the centrosome-equivalent spindle pole body.

In addition to dividing the genome, gamete formation also requires that gametes acquire organelles, either through inheritance prior to gamete membrane closure or through *de novo* synthesis (King et al., 2019; Otto et al., 2021; Sawyer et al., 2019; Suda et al., 2007). When gamete formation is complete, the lysosome-equivalent vacuole that originates from the mother precursor cell lyses and degrades all cellular content that was not inherited by the four gametes (Eastwood and Meneghini, 2015; Eastwood et al., 2013). In late meiosis, the resulting gametes continue to mature and then remain in a quiescent state inside a protective sack called the ascus (Neiman, 2005). When sufficient nutrients are detected, the gametes germinate and fuse with a gamete of opposite mating type, like egg and sperm fusion in metazoans, to reestablish the diploid state (Merlini et al., 2013; Rousseau et al., 1972).

Intuitively, the dramatic cellular changes that occur during gametogenesis must be coordinated at the level of gene expression, thus making yeast meiosis an excellent model for studying dynamic gene regulation in a natural context (Chu et al., 1998). Large-scale, multiomic sequencing efforts have revealed that over 90% of the yeast genome is actively translated at some point throughout gametogenesis with specific temporal fluctuations (Brar et al., 2012). In the majority of cases, the parallel measurements of transcription, translation, and protein levels in these studies revealed correlated patterns consistent with canonical gene regulation based on the central dogma (Cheng *et al*. 2018). However, these studies also provide evidence of unconventional transcript isoforms that are expressed at specific times throughout the meiotic program. Hundreds of extended transcripts have been identified in genome-wide sequencing efforts (Cheng et al., 2018; Chia et al., 2021; Tresenrider et al., 2021). This includes a subset of long undecoded transcript isoforms (LUTIs) that contain upstream ORFs (uORFs) that reduce translation efficiency of these transcripts (Chen et al., 2017; Chia et al., 2017). Additionally, hundreds of intragenic transcripts that potentially produce truncated protein isoforms have been identified, including an Ndt80-regulated intragenic *MRK1* transcript that plays a role in sporulation efficiency (Chia et al., 2021; Eisenberg et al., 2020; Zhou et al., 2017). While high levels of transcript heterogeneity has been well documented in both meiosis and mitosis, the biological impact of the majority of these gene isoforms remains unknown (Chia et al., 2021; Pelechano et al., 2013; Tresenrider et al., 2021).

Several genome-wide gain-of-function yeast libraries have been constructed previously (Arita et al., 2021; Douglas et al., 2012; Gelperin, 2005; Ho et al., 2009; Hu et al., 2007; Sopko et al., 2006). Although these libraries have been extremely valuable, each library overexpresses full-length open reading frames (ORFs) in the more conventional laboratory strain S288C, which exhibits low meiotic efficiency (Ben-Ari et al., 2006). Thus, systematic screening for functional meiotic gene isoforms has not been performed due to a lack of appropriate experimental tools. In this study, we use SK1 yeast, which undergoes meiosis with high efficiency, to construct five stage-specific inducible cDNA libraries that contain transcript isoforms that are expressed at a given meiotic stage. We use a computational approach to verify the representation of stage-specific genes in each library and confirm the presence of previously identified meiosis-specific gene isoforms. Finally, we develop a robust, multi-day competitive fitness screening and analysis pipeline to demonstrate the utility of our inducible cDNA libraries. This resulted in the identification of meiotic gene isoforms that are detrimental to competitive fitness when overexpressed, and we discovered a subset of mitochondrial proteins that cause dose-dependent disruption to respiration. Together, our study presents (i) a robust method for creating inducible cDNA libraries; (ii) meiosis-specific cDNA libraries for systematically studying meiotic gene and gene isoforms; and (iii) a dataset of meiotic genes and gene isoforms important for competitive fitness, which will require future study.

## RESULTS

### Isolation of phase-specific meiotic mRNA from synchronized yeast

Throughout the meiotic program, over 90% of the yeast genome is expressed, including meiosis-specific alternative transcript isoforms (Brar et al., 2012; Cheng et al., 2018; Chia et al., 2021; Tresenrider et al., 2021; Zhou et al., 2017). To construct inducible cDNA libraries that encompass the complete complement of meiotic isoforms, we set out to isolate mRNA from specific stages of meiosis. We utilized two previously established genetic systems that allow for highly synchronized progression of diploid yeast through the meiotic program. First, we used a strain containing a copper inducible promoter (*pCUP1*) upstream of two meiotic factors, *IME1* and *IME4,* that are required for meiotic entry (Berchowitz et al., 2013). Cells were grown in low nutrient sporulation media for 2 hours to arrest cells in a starvation state and then 50 mM of copper sulfate was added to the media to allow for synchronous progression through meiotic entry and S phase. The second strain contained a ß-estradiol-inducible system to turn on a chimeric *GAL4-ER* transcriptional activator driving *NDT80*, which is required for exit from pachytene of prophase I and allows for highly synchronized progression through Meiosis I and Meiosis II (Benjamin et al., 2003; Carlile and Amon, 2008). Cells were grown in sporulation media for 5 hours to achieve pachytene arrest and then 1 μM of ß-estradiol was added to the media to allow for synchronous progression through the meiotic divisions. Using these two strains, we performed parallel meiotic time courses and isolated mRNA over dense time points (Figure 1A). To ensure that our time courses were synchronized and reproducible, we performed each experiment in duplicate. Due to the fast nature of the meiotic divisions, spindle staining was used to count the percentage of cells in Meiosis I and Meiosis II in both replicates for the inducible *NDT80* system (Supplemental Figure 1A). Based on these results, we combined samples that corresponded to the following stages of meiosis: Starvation (pre-meiotic state), Early Meiosis (DNA replication and recombination), Meiosis I (first division), Meiosis II (second division), and Late Meiosis (gamete maturation). We next used mRNA-seq to assess the gene representation for each library. Using hierarchical clustering, we found that mRNA pools representing different meiotic stages had distinct gene expression profiles (Figure 1B) and mRNA pools representing different replicates from the same meiotic stage had a Spearman’s rank order correlation coefficient of over 0.9 for every library (Supplemental Figure 1B). Thus, our synchronized mRNA isolation protocol was reproducible and the resulting mRNA pools represent snapshots of five developmental phases of meiosis.

**Figure 1.**
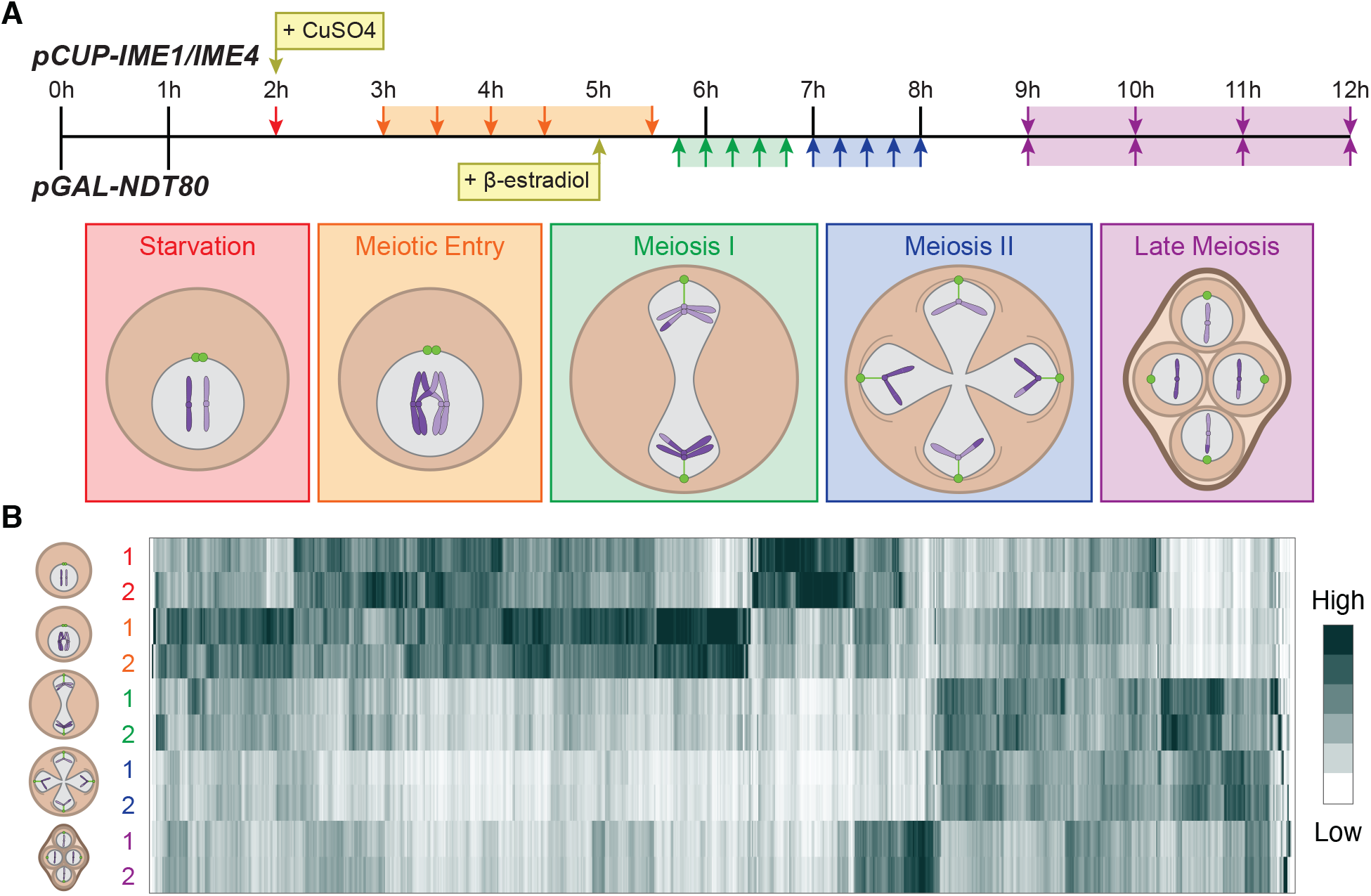
Meiotic synchronization allows for robust stage-specific isolation of mRNA. **(A)** A schematic outlining two parallel meiotic time courses using the indicated inducible genetic systems for cell synchrony. Arrows indicate when mRNA was harvested and samples with color-matched arrows were combined to produce mRNA libraries representing five meiotic stage. **(B)** Heat map displaying relative mRNA expression profiles from (A) ordered by hierarchical clustering. Two replicates for each time course are shown.

### Construction of stage-specific, inducible meiotic cDNA library

Using our stage-specific mRNA pools, we next wanted to construct inducible plasmid libraries that could be expressed in yeast (Figure 2A). Due to the high-level of reproducibility in our two meiotic time courses, we decided to proceed with mRNA isolated from “Replicate 1” for all libraries (Figure 1). First, we synthesized complementary DNA (cDNA) from each of our mRNA pools using the highly processive TGIRT-III enzyme, which has been shown to outperform other commercially available reverse transcriptases (Mohr et al., 2013). After converting each meiotic mRNA pool to cDNA, we inserted the libraries into a Gateway donor vector (pDON222) and amplified the libraries in *Escherichia coli* (Katzen, 2007). We next built a high copy number destination vector (pUB914) containing the *GAL10* promoter, which would allow for galactose induction of our cDNA libraries. The cDNA in the donor vector was moved to this destination vector and again amplified in *E. coli.* Finally, we built an SK1 strain that would be amenable to a pooled screening workflow. Unlike other laboratory yeast strains such as S288C or W303, wild-type SK1 flocculates and has poor mother-daughter separation following cytokinesis, which causes cells to clump together. Additionally, SK1 has a defective *GAL3* gene that is critical for galactose-based induction. Thus, we deleted *FLO8* to remove the ability of cells to flocculate, replaced the SK1-specific allele of *AMN1* with a S288C-specific allele to promote mother-daughter cell separation following cytokinesis, and replaced the defective SK1 *GAL3* gene with a functional W303 *GAL3* gene. We then transformed each of the five inducible meiotic cDNA libraries into this new SK1 screening strain using electroporation.

**Figure 2.**
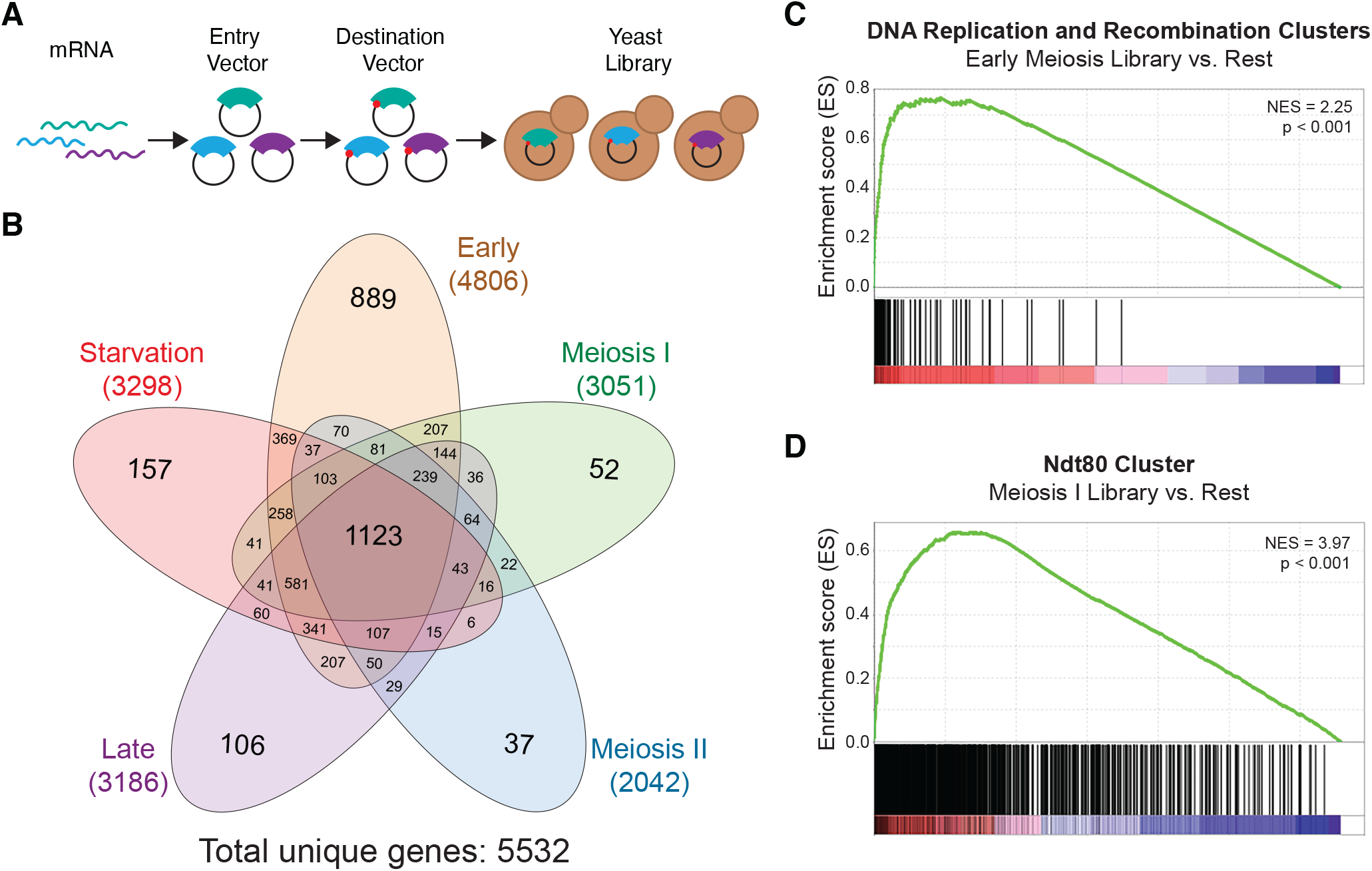
Gene representation in meiotic cDNA libraries. **(A)** A schematic of inducible yeast cDNA library construction. TGIRT reverse transcriptase was used for cDNA synthesis and Gateway cloning was used for plasmid cloning. Plasmids were transformed into yeast using electroporation. **(B)** DNA-seq was used to estimate the number of genes represented in the indicated libraries. The Venn Diagram shows cDNAs that are unique or common between the 5 libraries. **(C)** Gene set enrichment analysis (GSEA) of all stages of library construction for the “Early Meiosis” library versus the other four libraries. Vertical black bars represent positioning of genes in the DNA replication and recombination clusters from Brar et al., 2012. The heat map indicates genes that are more enriched in the “Early Meiosis” libraries and blue indicates genes that are more enriched in the other libraries. NES, Normalized Enrichment Score. **(D)** GSEA analysis as in (C), except all stages of library construction for the “Meiosis I” library is being compared to the other four libraries. Vertical black bars represent positioning of genes in the Ndt80 cluster from Cheng et al., 2018. The heat map indicates genes that are more enriched in the “Meiosis I” libraries and blue indicates genes that are more enriched in the other libraries. NES, Normalized Enrichment Score.

To define the number of genes that were present in each of the five meiotic cDNA libraries we used DNA-seq to determine the representation of genes in each library relative to the initial mRNA pools at each step of construction: after insertion into the entry vector, after insertion into the destination vector, and after transformation into yeast (Supplemental Table 1). To reduce the amount of vector backbone sequences, the cDNAs were PCR amplified prior to DNA sequencing library preparation and then sequenced on an Illumina platform. Sequencing reads were aligned to the SK1 genome using HISAT2 and transcripts per million reads (TPM) for each gene was calculated using STRINGTIE (Pertea et al., 2016). It is important to note that TPM was developed to measure RNA expression levels, but in this study, we use TPM to quantify the abundance of a particular cDNA. To eliminate background noise, we reasoned that genes represented in each library should have a TPM count greater than 5 at each step of library construction (mRNA, entry vector, destination vector, yeast) (Supplemental Figure 2A; Supplemental Table 2). Using these criteria, we found that our libraries contained 5563 unique genes, which is more than 84% of the yeast genome (Figure 2B). Each library contained a unique set of genes and we found 1123 genes that were common to all 5 libraries, which we term “housekeeping genes” since these genes are kept on to some extent during starvation state and for the entire duration of meiosis. Consistent with this nomenclature, gene ontology (GO) enrichment analysis revealed housekeeping genes were enriched for genes involved in catabolism and proteolysis (Supplemental Table 2; Raudvere et al., 2019).

As a quality control check, we performed gene set enrichment analysis (GSEA) to ensure that stage-specific genes were being enriched in the appropriate libraries (Mootha et al., 2003; Subramanian et al., 2005). First, we compared the DNA-seq data for all stages of the “Early Meiosis” library construction (mRNA, entry vector, destination vector, and yeast) against DNA-seq data for all stages of library construction for the rest of the four cDNA libraries (termed “Rest” in Figure 2C; Supplementary Table 3). We defined a set of genes that we expected to be expressed in early meiosis by combining the “DNA replication cluster” and “recombination and SC formation cluster” from a previous study (Brar et al., 2012). We found a highly significant enrichment of these genes in the “Early Meiosis” libraries, suggesting that release of cells into early meiosis using the *pCUP-IME1/pCUP-IME4* system was highly synchronized. Next, we repeated the GSEA analysis but instead compared all stages of the “Meiosis I” library construction against all stages of library construction for the “Rest” of the four remaining libraries (Figure 2D; Supplementary Table 3). In this case, we defined a set of genes that come on at the same time as the key meiotic regulator, Ndt80, which was termed the “Ndt80 cluster” in a previous study (Cheng et al., 2018). Again, we found highly significant enrichment of genes that come on concomitant with Ndt80 in the Meiosis I library suggesting that release of cells into Meiosis I using the *pGAL-NDT80* system was also highly synchronized. Thus, we conclude that the meiotic time courses were precise, and the library construction pipeline maintains accurate gene representation in all cDNA libraries.

Lastly, we wanted to determine whether any size-dependent bias was introduced into our libraries throughout the library construction process (Supplemental Figure 2B; Supplemental Table 4). We plotted the size distribution of the yeast genome, which contains 6574 genes, and found an average gene size of 1351 bp and a median gene size of 1076 bp. Compared to the 5563 genes found in our cDNA libraries, we found an average gene size of 1438 bp and a median gene size of 1181 bp suggesting that a subset of small genes may have been lost during library construction. We suspected that genes smaller than 200 bp may have been lost at a DNA column purification step (see Methods), but examination of the size distribution of the 1011 genes missing from our cDNA libraries reveal an average gene size of 870 bp and a median gene size of 446 bp suggesting that many missing genes must be absent due to other reasons. Gene ontology (GO) enrichment analysis revealed many genes involved in ribosome biogenesis and RNA metabolism are missing from the cDNA libraries, which are two processes known to be down regulated during meiosis (Supplemental Table 4; Raudvere et al., 2019). Thus, we suspect that genes that were lost in library construction are likely genes that are not expressed or expressed at very low levels during gametogenesis. Comparison of gene size distribution between the five libraries shows a similar pattern despite a difference in total genes in each library suggesting that library construction was consistent between libraries (Supplemental Figure 2C). Thus, we find that our meiotic cDNA library construction pipeline retains a high level of gene representation for each stage of meiosis.

### Meiotic cDNA libraries contain meiosis-specific mRNA transcripts

In addition to estimating the number of genes in our cDNA libraries, we also wanted to determine whether meiosis-specific transcript isoforms were present. Recent genome-wide studies have identified hundreds of alternative transcription start sites (TSSs) and transcription end sites (TESs) during the meiotic program (Chia et al., 2021; Tresenrider et al., 2021; Zhou et al., 2017), although the functional relevance of less than a handful is known (Chen et al., 2017; Chia et al., 2017; Tang et al., 2004; Zhou et al., 2017). We decided to focus on identifying shorter isoforms of annotated genes in our libraries because they could theoretically produce non-canonical truncated proteins with altered cellular function. We first set out to identify cDNAs in our destination vector libraries that were truncated at the 5’-end, which could represent intragenic isoforms. We reasoned that 5’-truncated cDNAs could originate from two sources: either from technical artifacts associated with reverse transcription or true meiosis-specific mRNAs. We started by assessing the processivity of the TGIRT-III reverse transcriptase by determining if any size-dependent 3’-bias for read density was detectable in our DNA-seq data. Since the processivity of the TGIRT-III reverse transcriptase is proportional to the size of a gene, we predicted that 3’-bias, if it existed, would be highly detectable in larger genes and undetectable in smaller genes. Thus, we divided the entire yeast genome into 4 equal quartiles by gene size: small (100 – 577 bp), medium (578 – 1087 bp), large (1088 – 1773 bp) and extra-large (1774 – 17283 bp). For each library, we took any gene with a TPM > 100 in the destination vector pool, divided the gene into 10 equal segments, and quantified the TPM per segment, which was then normalized by dividing each segment by the maximum TPM across the 10 segments for a given gene. We then plotted the average normalized TPM values across the 10 segments for all genes in each size quartile (Figure 3A). We did not observe any general size-dependent 3’-bias in read density consistent with the TGIRT-III reverse transcriptase being highly processive (Mohr et al., 2013).

**Figure 3.**
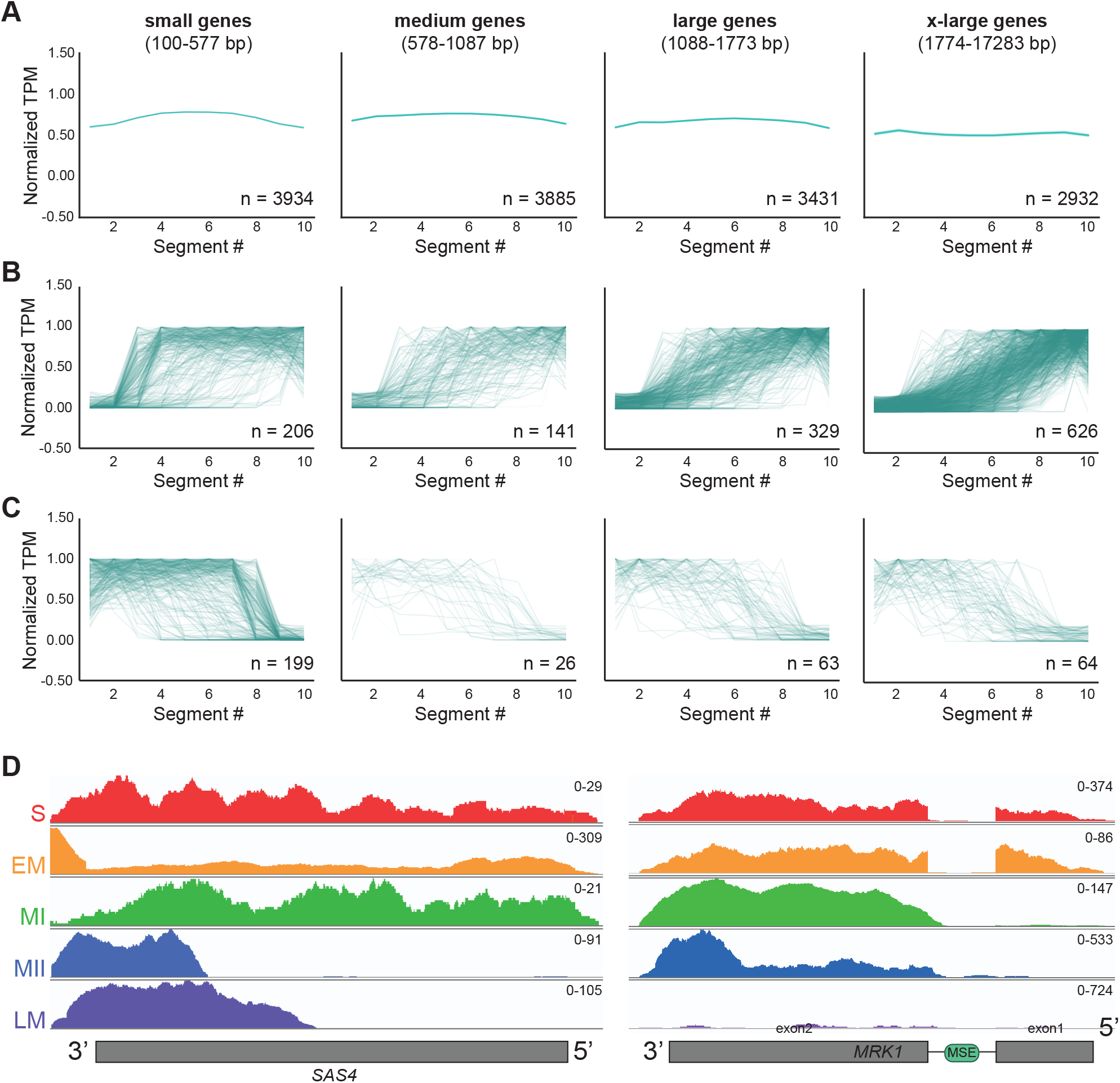
Identification of truncated gene isoforms in meiotic cDNA libraries. **(A)** cDNAs with the indicated size range from all libraries were divided into 10 equal segments and the average normalized TPM (transcripts per million reads) per segment was plotted. **(B)** cDNAs from any library that had low TPM in segments 1 and 2 and high TPM in segments 9 and 10 were considered potential 5’-truncations. The normalized TPM per segment are plotted for all 5’-truncations in a given size quartile. **(C)** cDNAs from any library that had high TPM in segments 1 and 2 and low TPM in segments 9 and 10 were considered potential 3’-truncations. The normalized TPM per segment are plotted for all 5’-truncations in a given size quartile. **(D)** Read density plot for *SAS4* across 5 yeast cDNA libraries, which was previously identified as a having a truncated transcript and corresponding protein isoform in meiosis (Brar et al., 2012; Chia et al., 2021). The y-axis scale for number of reads is found a the top right corner of each plot. S, Starvation; EM, Early Meiosis; MI, Meiosis I; MII, Meiosis II; LM, Late Meiosis. **(E)** Read density plot for *MRK1* across the 5 yeast cDNA libraries (as in (D)), which was previously identified as having a truncated transcript isoform important for meiotic progression (Zhou et al., 2017).

To determine if 3’-bias was seen in the largest genes in our cDNA libraries, we pulled the top 3 largest genes that showed strong evidence of 3’ bias for further examination. To do this, the largest genes that had the very low TPM for segments 1-3 (average TPM for segments 1-3 less than 1.5) and the maximum TPM for segment 10 (TPM greater than 20) (Supplemental Figure 3A). Altogether, this analysis pulled 11 unique extra-large genes with potential 3’ bias that were all over 5,600 bp in length. Interestingly, each of these extra-large genes showed a steep drop in read density after approximately 2,000 bp. Although we cannot rule out that these reads originated from *bono fide* meiotic truncations, we conclude that truncations found in genes greater than 2,000 bp in size may be a technical artifact and so should be verified by cross-referencing to other genome-wide TSS studies (Chia et al., 2021). Since 4356/5563 (over 78%) genes in our cDNA libraries are less than 2000 bp, we conclude that the TGIRT reverse transcriptase introduces minimal 3’-bias into our libraries with only a subset of extra-large genes being potentially affected.

We next wanted to estimate how many meiotic transcript isoforms exist in each library. Genes that had more reads at the 5’-end (segments 1 and 2) of the gene versus the 3’-end (segments 9 and 10) were identified and considered to be genes with potential 5’-truncated isoforms (Figure 3B; Supplemental Table 5). In total, we identified 1302 genes with potential 5’-truncations, with 48% being found in extra-large genes. A recent study used multiple sequencing-based techniques to define a comprehensive set of genes that express alternative transcript isoforms throughout the meiotic program (Chia et al., 2021). Cross referencing with TSSs identified in this study confirmed 208/1302 genes (16%) that had detectable intragenic TSSs. Included in this list was a transcript isoform shown to make a corresponding truncated protein *SAS4* (Figure 3D). Additionally, we identified the intergenic transcript of *MRK1*, which was previously found to be regulated by Ndt80 and be important for efficient meiotic progression (Figure 3E; Zhou et al., 2017). It is important to note that visual inspection of a subset of other 5’-truncations identified by our computational approach revealed a high rate of false positives due to reads from adjacent or overlapping genes, thus we will continue referring to these subsets as 5’-truncation candidates.

To gain a sense of how sensitive our computational pipeline was for identifying genes with intragenic start sites, we manually checked the 23 genes identified in Zhou et al. (2017) that were confirmed by tiling array transcription dataset from another study (Lardenois et al., 2011). Visual examination of read density of our yeast cDNA libraries provides evidence that 13/24 of these 5’-truncated genes were present in our libraries, however, only five of these genes were identified by our computational pipeline (Supplemental Figure 3B). We cross referenced with the TSSs in Chia et al. (2021) and found that five of these genes were not found in either of our computational analyses. Further inspection revealed that many of these genes had lower TPM counts in segment 10, causing these genes to fall under the threshold for detection (see Methods). Whether these truncated isoforms are biologically relevant, or a technical artifact remains unknown. Taken together, our computational pipeline can recapitulate previously identified intragenic transcripts within our cDNA libraries, however there is a high false positive rate and limitations as to what read density patterns can be robustly identified.

We also wanted to identify genes that had more reads at the 3’-end of the gene versus the 5’ end and considered these genes to have potential 3’-truncated isoforms. Using the reciprocal analysis for the 5’-truncations, we identified 351 genes that had potential 3’-truncated isoforms, and 78 had detectable internal transcription end sites (TESs) in Chia et al. (2021) (Figure 3C; Supplemental Table 5). Although the analysis pipeline for the 3’-truncation candidates has similar caveats to the 5’-truncation pipeline, we conclude that a subset of genes with 3’-truncated isoforms exist in our cDNA libraries.

### Systematic examination of meiotic gene overexpression on competitive fitness

We next developed a competitive fitness assay to identify all genes or gene isoforms that are beneficial or detrimental for decrease competitive cell growth (Figure 4A). To begin the screen, a frozen aliquot of each yeast library was grown to logarithmic phase in uninducing (media containing raffinose) media for 8 hours. Next, each library was split into uninducing (media containing glucose; YPD) or inducing (media containing galactose; YPGR) media and allowed to grow to saturation over 24 hours. Each day, a sample of each saturated culture was harvested for DNA sequencing and each culture was then diluted 1 in 1,000 in the same corresponding media for another day of growth. This cycle was repeated over 5 days and optical density (OD_600_) measurements revealed that each day represented approximately 8 generations of growth for the entire library. Each screen was performed in triplicate and plasmids from each sample were isolated, prepared for DNA-seq, and analyzed as in Figure 2.

**Figure 4.**
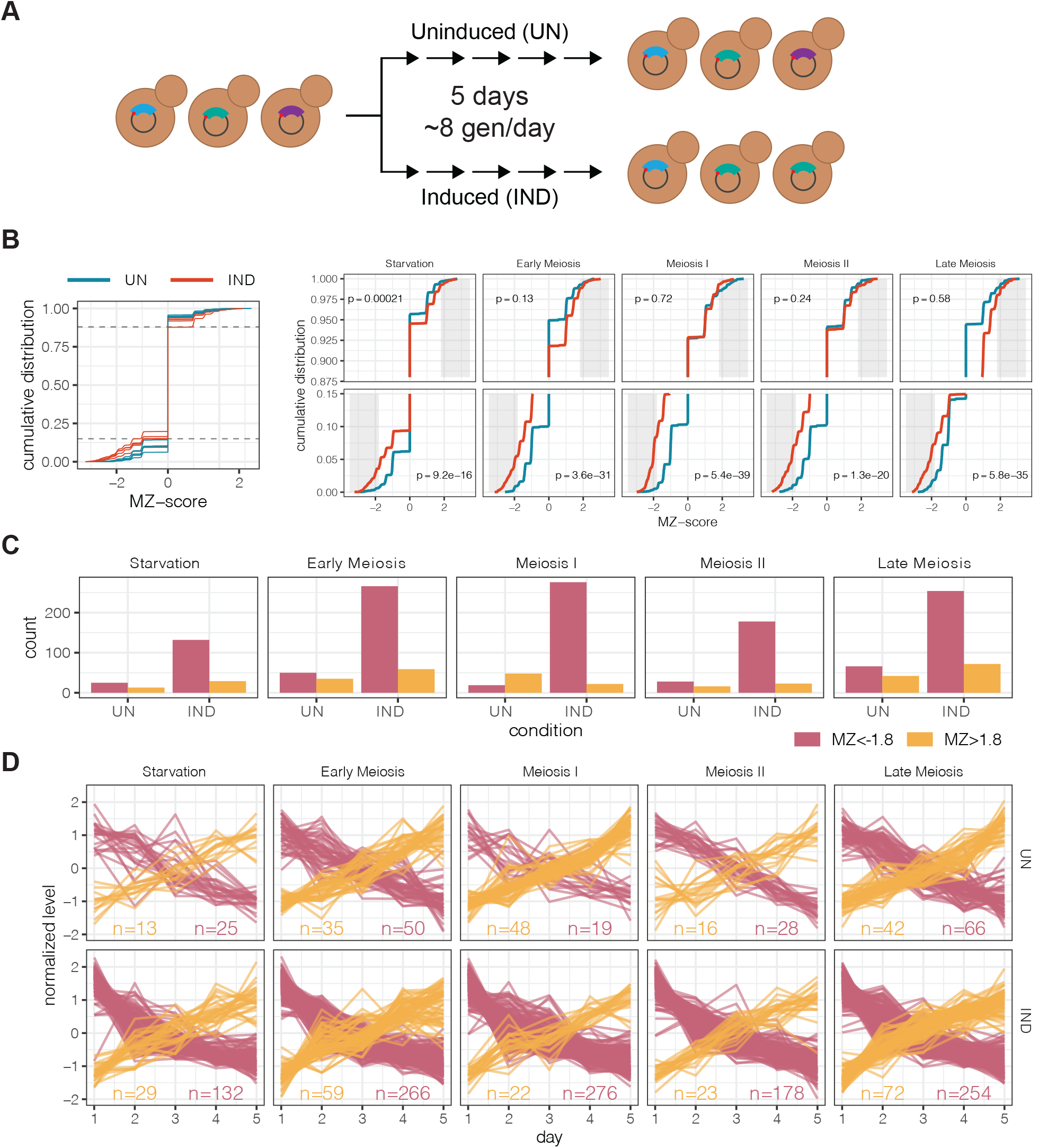
Assessing the effect of meiotic gene expression on competitive fitness. **(A)** Schematic of competitive assay. Each meiotic library was grown in uninducing (UN) or inducing (IND) conditions for 5 days. Cultures were grown to saturation each day, a sample was taken for DNA-seq, and the culture was re-diluted 1 in 1000 for another day of growth (∼ 8 divisions per day). **(B)** Monotonicity Z-scores (MZ-scores) were calculated for every gene in uninducing (UN) and inducing (IND) conditions and the cumulative distribution is shown (left plot). Zoomed-in plots for data above and below the dotted lines are shown for the individual libraries on the right. Grey boxes highlight MZ-score of −1.8 (top row) and 1.8 (bottom row), which is the cut-off for genes considered to be disenriched or enriched over time, respectively. P-values signify whether induction of the cDNA library causes a detectable change in the number of genes becoming disenriched (top row) or enriched (bottom row). **(D)** The number of genes that are disenriched (red) or enriched (yellow) in uninducing (UN) or inducing (IND) conditions. **(E)** Normalized levels of genes that disenrich (red) or enrich (yellow) over the course of 5 days are plotted for uninducing (top row) or inducing (bottom row) conditions.

In order to identify cDNAs that enriched or disenriched in the population throughout the competitive fitness screen, we used monotonicity z-scores (MZ-scores) to identify genes that showed incrementally increasing or decreasing TPM over time (adapted from Taliaferro et al., 2016; Supplemental Table 6). We found that induction of only the pre-meiotic “Starvation” cDNA library caused a significant change in the number of genes that were beneficial for competitive growth (MZ-score > 1.8); however, we observed that induction of all cDNA libraries caused a significant increase in the number of genes that are detrimental for competitive growth (MZ-score < −1.8) (Figure 4B). Each library resulted in different numbers of genes that affect competitive fitness (Figure C-D). In total, we found 188 genes that were beneficial for competitive fitness and 877 genes that were detrimental for competitive fitness in at least one screen (Supplemental Figure 4A-B). Interestingly, we identified 6 “housekeeping genes” that were detrimental, including *TIM23*, *SDH4*, *CCC1*, and *YPT1*; four genes that have been previously shown to cause dosage sickness or lethality (Douglas et al., 2012; Sopko et al., 2006; Yoshikawa et al., 2011).

We were interested in determining whether any of the genes that showed negative effects on competitive fitness represented meiosis-specific isoforms. We reasoned that induction of gene isoforms that facilitated a meiosis-specific function would be detrimental to competitive fitness. We overlapped detrimental gene datasets with the 5’- and 3’- truncation lists produced in Figure 3B-C and found 71 genes that had potential 5’-truncations and 39 genes that had potential 3’- truncations that were detrimental for competitive fitness (Supplemental Table 7). Taken together, these proof-of-principle screens reveal that multi-day growth of the meiotic cDNA library can select for cDNAs that are beneficial or detrimental to competitive fitness. Furthermore, our cDNA libraries identify a subset of gene isoform candidates that affect competitive fitness, which will require future study.

### Comparing toxic genes from meiotic libraries to other gain-of-function screens

We next wanted to ensure that the proof-of-principle assay using our inducible, meiotic cDNA libraries was capable of recapitulating previously published data produced by other gain-of-function screens. Previously, two other studies have performed competitive assays with pooled, genome-wide gain-of-function libraries in budding yeast (Arita et al., 2021; Douglas et al., 2012). Rather than creating cDNA libraries from mRNA pools, both groups systematically constructed arrayed overexpression libraries in the S288C background, which were then pooled. In Douglas et al. (2012), the authors pooled an arrayed library containing a low copy number plasmid (CEN/ARS) that overexpressed a given gene using a galactose-inducible promoter (GAL1/10). The pooled library was grown in inducing conditions in continuous logarithmic phase for 20 generations resulting in identification of 361 toxic genes. In Arita et al. (2021), the endogenous promoter of each gene was replaced with a synthetic promoter that can be activated by a β−estradiol-inducible engineered transcription factor. Competitive fitness assays were performed for 36 hours in synthetic complete (SC) media or 48 hours in yeast nitrogen base (YNB) media resulting in the identification of 432 genes that are detrimental for competitive growth in at least one condition.

We found that 15.4% of dosage lethal genes overlapped with these two previous studies (8.8% with Arita et al. (2021) and 5.6% with Douglas et al. (2012)) while the overlap between Arita et al. (2021) and Douglas et al. (2012) was over 20%. We predicted that this low overlap could be due to technical differences between the assays: (i) our screen was performed in SK1 instead of BY; (ii) our cDNA libraries are expressed using a galactose-inducible promoter on a high copy number plasmid, which would result in much higher expression levels than the other two studies; (iii) our time course was longer; and (iv) our cultures were not maintained in the logarithmic state. To gain a better understanding of what genes were identified in our study, we performed gene ontology (GO) enrichment on the 760 genes not identified in the other two studies and observed an abundance of mitochondrial-related GO terms (Figure 5B) (Raudvere et al., 2019). Cross-referencing with the *Saccharomyces* Genome Database (https://www.yeastgenome.org/) showed that 251/760 of our unique hits had functions related to the mitochondria (GO:0005739; Supplemental Table 8). To determine how many of these genes produced proteins that localized to the mitochondria, we wanted to determine the number of genes that contained a mitochondrial targeting sequence (MTS). We created a list of genes containing a predicted MTS according to four published algorithms (Almagro Armenteros et al., 2019; Bannai et al., 2002; Claros and Vincens, 1996; Fukasawa et al., 2015) and found that 43 of our mitochondria-related hits contained an MTS (Supplemental Table 8). Thus, competitive screening of our cDNA libraries could recapitulate results from previous screens and lead to the identification of mitochondrial proteins important for competitive fitness.

**Figure 5.**
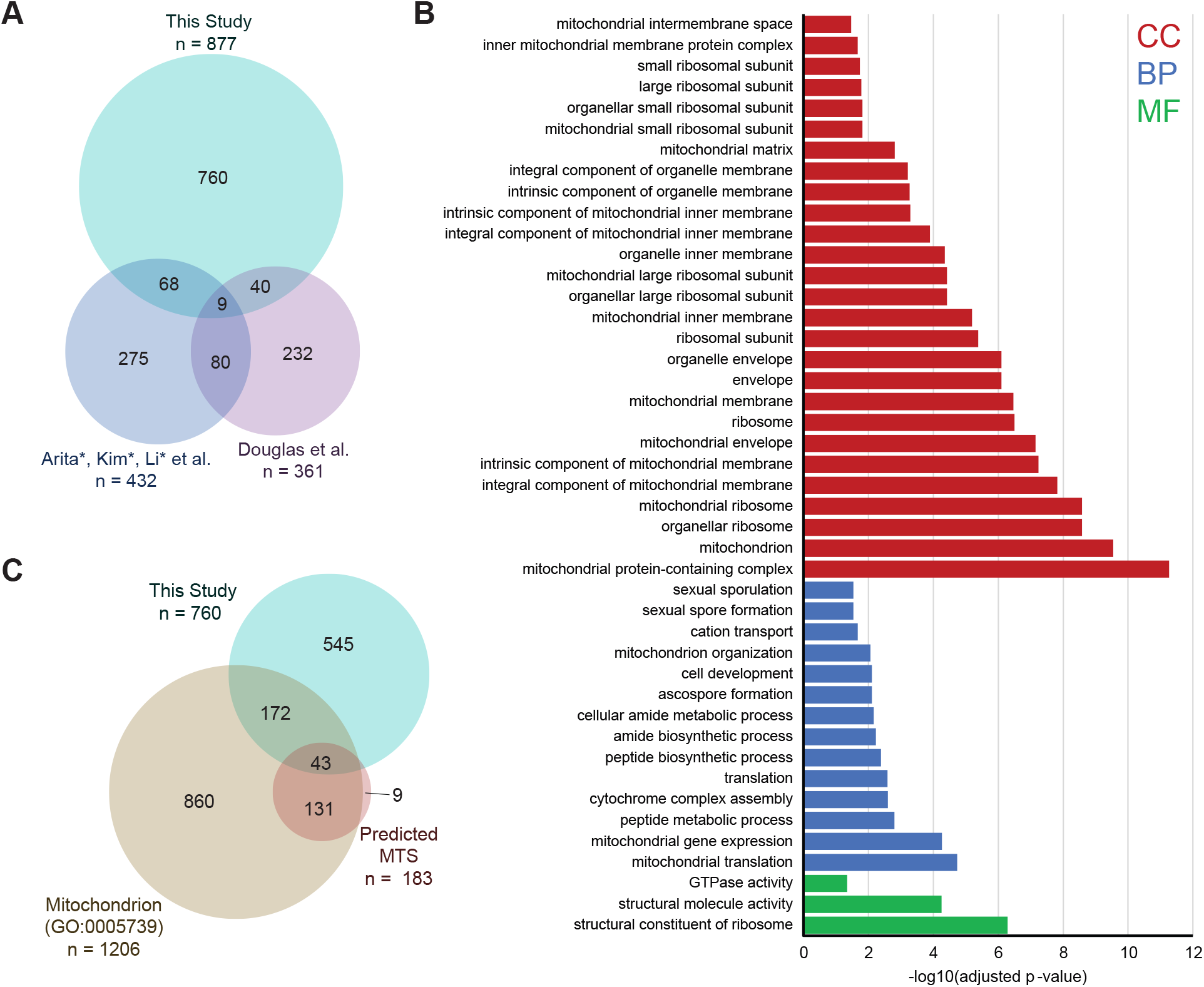
Overexpression of meiotic cDNA libraries reveals a subset of mitochondrial genes that are detrimental to competitive fitness. **(A)** Overlap between genes identified in this study (SK1 background) to be detrimental for competitive fitness and genes identified in other studies (BY background) using a similar assay (Arita et al., 2021; Douglas et al., 2012). **(B)** Gene ontology (GO) enrichment analysis of the 760 genes found to be unique to this study in (A). Only GO terms that contain less than 1000 genes is shown. CC, cellular component; BP, biological process; MF, molecular function. **(C)** Of the 877 genes found to be detrimental for competitive fitness, 254 are annotated as being mitochondria-related and 50 have a predicted mitochondrion targeting sequence (MTS).

### Overexpression of mitoproteins causes reduced respiration

In standard yeast growth conditions, cells will grow exponentially when nutrients are in excess. As a culture becomes more densely populated, the yeast will transition into a saturation phase where growth rate will slow and eventually stop due to nutrient deprivation and over-crowding. During this saturation phase, yeast cells will undergo a diauxic shift from fermentation to respiration, a form of metabolism that is more dependent on functional mitochondria. Because our 5-day competitive assay allowed cultures to become saturated each day prior to re-dilution, we hypothesized that the disenrichment of mitochondrial genes in our competitive assays was due to defects in cellular respiration. To test this, we reconstructed overexpression plasmids of 9 mitochondrial genes (*ALD4*, *PSD1*, *COX2*, *TIM44*, *MPS9*, *ATP16*, *MIA40*, *ABF2*, *COX11*) that became disenriched during our competitive fitness screens and contained a predicted MTS for further growth analyses.

First, we wanted to confirm that overexpression of these mitochondrial genes caused a competitive fitness defect. To measure this, we used flow cytometry to measure competitive fitness between cells overexpressing each mitochondrial gene against cells containing an empty vector. We created two strains: one containing a *PGK1-GFP* allele marked by a *HIS3* cassette (called the “GFP+” strain) and an untagged strain containing *HIS3* at the native locus (called the “GFP-” strain) to match the auxotrophy of the GFP+ strain. Overexpression plasmids containing the indicated mitochondrial gene were transformed into each strain background and competitive assays were performed in two orientations. First, GFP- strains overexpressing a mitochondrial gene were mixed 50:50 with a GFP+ strain containing an empty vector (Figure 6A). Initially, after competitive cultures grow in inducing conditions for one day, we see approximately even representation of the GFP- and GFP+ strains, suggesting that mitochondria were competent in both strains. However, we see that in all cases, except for the empty vector: empty vector control, the GFP+ empty vector strain outcompetes the GFP- strain after two days of growth in inducing conditions suggesting competitive fitness defects in the GFP- strain overexpressing the mitochondrial gene. Second, the reciprocal experiment was performed where GFP+ strains overexpressing a mitochondrial gene were mixed 50:50 with a GFP- strain containing the empty vector (Figure 6B). Consistent with the previous experiment, the GFP- empty vector strain outcompeted all GFP+ strains overexpressing a mitochondrial gene after two days of growth in inducing conditions. Thus, these experiments reveal that overexpression of each of these mitochondrial genes negatively affects competitive fitness.

**Figure 6.**
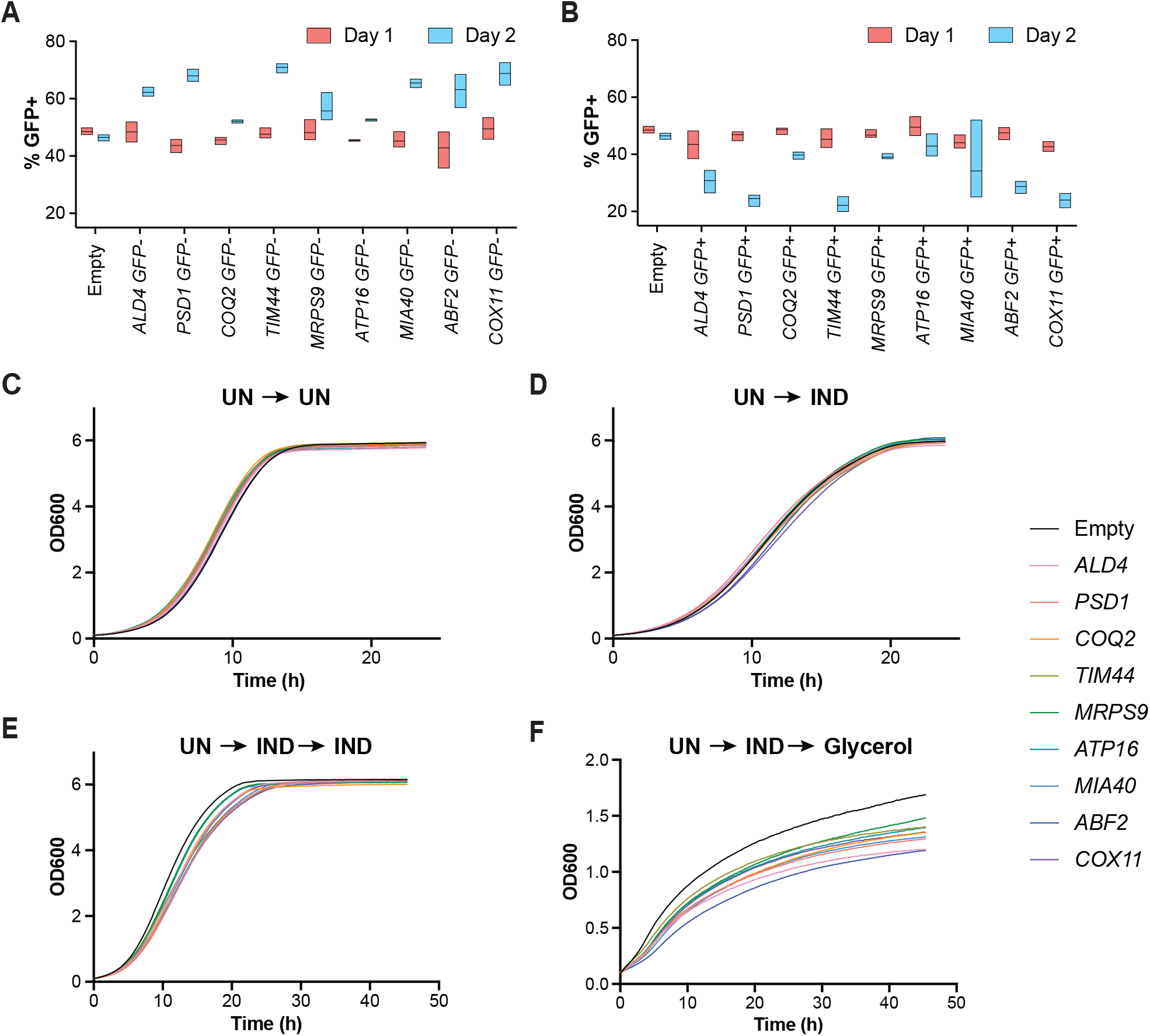
Overexpression of mitochondrial proteins causes disrupts respiration. **(A)** Strains over expressing the indicated genes and untagged *PGK1* were grown in competition with an equal number of cells containing an empty vector and *PGK1-GFP*. The percentage of GFP+ strain by flow cytometry is plotted after 1 and 2 days of growth in inducing media. Data for 3 biological replicates are shown. **(B)** Strains over expressing the indicated genes an *PGK1-GFP* were grown in competition with an equal number of cells containing an empty vector and untagged *PGK1.* The percentage of GFP+ strain by flow cytometry is plotted after 1 and 2 days of growth in inducing media. Data for 3 biological replicates are shown. **(C)** Plasmids containing the indicated genes were grown in uninducing conditions overnight and then diluted into uninducing media for growth measurements over 24 hours. Each growth curve represents the average of 3 biological replicates. **(D)** Growth curves as in (C) except cells were grown in uninducing conditions overnight and then diluted into inducing media for growth measurements over 24 hours. Each growth curve represents the average of 3 biological replicates. **(E)** Growth curves as in (C) except cells were grown in uninducing conditions overnight, then diluted into inducing media for 24 hours, then diluted into inducing media for growth measurements over 48 hours. Each growth curve represents the average of 3 biological replicates. **(F)** Growth curves as in (C) except cells were grown in uninducing conditions overnight, then diluted into inducing media for 24 hours, then diluted into non-fermentable media (glycerol) for growth measurements over 48 hours. Each growth curve represents the average of 3 biological replicates.

We next wanted to test whether overexpression of these mitochondrial proteins had any specific effect on cellular respiration. First, we recovered each strain in uninducing conditions overnight and then independently diluted cultures into inducing or uninducing media and measured growth over 24 hours (Figure 6C-D). We did not observe differences in growth rate between the mutant strains and the empty vector control in either condition. Next, we tested whether any defects in respiration were detectable after mitochondrial genes were overexpressed for 24 hours (Figure 6E-F). We allowed each strain to grow in inducing media for 24 hours and then diluted cells into inducing media or media containing glycerol, a non-fermentable carbon source that requires respiration to be metabolized. We found modest defects in growth rate as cells grew in inducing media for a second day but observed a large reduction in growth rate in glycerol. Together, these results suggest that overexpression of mitochondrial genes has no effect on non-competitive growth but does have a negative effect on cellular respiration, which is important for competitive fitness during the saturation phase of growth.

## DISCUSSION

We have constructed five stage-specific, inducible cDNA libraries from meiotic mRNA and performed a multi-day proof-of-principle screen to demonstrate its utility. We anticipate that this study will be useful in multiple ways. First, our comprehensive gain-of-function dataset provides new information about meiotic genes and gene isoforms that are beneficial or detrimental to competitive fitness in mitotically growing cells. To date, hundreds of alternative TSSs and TESs have been detected, but the function of most of these alternative isoforms remains unknown. Our competitive assay revealed 877 genes that are detrimental to competitive fitness, including 110 potential transcript isoforms. If we assume that functionally relevant meiotic transcripts may be detrimental for mitotic growth, further analysis of these meiotic genes and gene isoforms may reveal novel biological insight into how gametogenesis is orchestrated.

The meiotic cDNA libraries and SK1 screening strain from this study can be used as a tool to study many other aspects of gametogenesis. Studies in budding yeast reveal that gametogenesis naturally eliminates age-associated damage from gametes, which leads to lifespan resetting (Unal et al., 2011). Identifying the cellular pathways that must converge to completely rejuvenate gametes remains an active area of research. In particular, the cDNA libraries could be used in gain-of-function screens in mitotic cells to identify regulators of meiosis-specific gene expression, organelle remodeling, and quality control pathways (Neiman, 2005; Otto et al., 2021; Sawyer et al., 2019; Suda et al., 2007). Furthermore, gametogenesis has been shown to eliminate many types of age-associated senescence factors including excess nucleolar material, protein aggregates, and non-chromosomal ribosomal DNA (rDNA) circles (King et al., 2019; King and Ünal 2020). Thus, our meiotic cDNA libraries provide an excellent tool for studying natural rejuvenation pathways as well as for determining whether these rejuvenation pathways can be leveraged to counteract cellular aging in mitotically growing cells.

Our multi-day time course allowed yeast cDNA libraries to reach saturation daily before being re-diluted. As cells increase in density and run out of nutrients, the cells must undergo a diauxic shift to transition to respiratory growth, a process that relies on mitochondrial function. This feature of our screen resulted in the identification of 43 genes containing a mitochondrial targeting sequence (MTS) that are important for competitive fitness and were not previously identified in other competitive screens (Arita et al., 2021; Douglas et al., 2012). Recently, mis-localized mitochondrial proteins have been found to elicit multiple branches of the mitoprotein stress response and disrupt proteostasis (Boos et al., 2020). Our screening pipeline inadvertently produced a list of mitochondrial proteins that become dosage-lethal during competitive growth and we find that overexpression of at least a subset of these proteins cause defects in cellular respiration. How overexpression of these mitoproteins disrupt respiration remains an open question for future studies.

Lastly, we developed a robust cDNA library construction pipeline and a simple, multi-day competitive fitness assay that could be applied to study other biological context where alternative transcript isoforms are thought to play a role. Studies in yeast and human cells have shown that, similar to gametogenesis, changes in stress or nutrients causes expression of alternative transcripts to modulate gene expression changes (Hollerer et al., 2016; Pelechano et al., 2013). Thus, the construction pipeline to create unbiased cDNA libraries presented in this study may provide a fast and easy strategy for assessing the effect of overexpressing non-meiotic gene isoforms. Additionally, alternative transcript isoforms have been shown to be important in various aspects of development, health, and disease in a variety of organisms (Baillie et al., 2014; Demircioğlu et al., 2019; Reyes and Huber, 2018; Xia et al., 2014; Yang et al., 2020). Thus, our computational analysis pipelines may also be useful in identifying gene isoform candidates in both yeast and higher eukaryotes to gain insight about these processes.

In summary, this study provides a robust method for creating inducible cDNA libraries for screening in budding yeast. We have focused our efforts on creating five galactose-inducible meiotic cDNA libraries in high copy number plasmids, which allowed us to create computational methods to detect transcript isoforms. Furthermore, we developed a competitive fitness-based screening pipeline amenable to our cDNA library and others, which revealed a subset of mitochondrial proteins that are important for cellular respiration. The data from this study will help to prioritize future studies of meiotic transcript isoforms, and we anticipate that these libraries, as well as similar cDNA libraries that may be built in the future, will facilitate a more systematic interrogation of alternative transcripts at the cellular level in different developmental and/or environmental conditions.

## MATERIALS AND METHODS

### Yeast strains, plasmids, and primers

All strains in this study were derived from the SK1 background. Strain genotypes, plasmids used for strain construction, and primer sequences used in this study can be found in Supplemental Table 9. Individual yeast strains were thawed from the −80°C freezer onto YPG plates (1% yeast extract, 2% peptone, 3% glycerol, 2% agar) overnight at 30°C to select for cells with healthy mitochondria. Haploid cells were then plated on YPD (1% yeast extract, 2% peptone, 2% glucose, 2% agar) plates or diploid cells were plated on YPD with high glucose (1% yeast extract, 2% peptone, 4% glucose, 2% agar) to prevent sporulation and incubated at 30°C overnight prior to experimentation. For competitive fitness screens, one tube of the yeast cDNA library was freshly thawed and used for direct inoculation of a liquid culture. cDNA libraries were never refrozen. All yeast grown in liquid media were incubated at 30°C while shaking.

### Meiotic time courses

Messenger RNA was isolated from synchronized meiosis time courses using strains A33366/UB1007 and A22678/UB140. For both strains, cells were grown in liquid YPD media (1% yeast extract, 2% peptone, 2% glucose, 22.4 mg/L uracil, and 80 mg/L tryptophan) overnight at 30°C. Cultures were then diluted to OD_600_ = 0.25 in BYTA media (1% yeast extract, 2% bacto tryptone, 1% potassium acetate, and 50 mM potassium phthalate) and grown for 15 hr at 30°C. When OD_600_ was over 5, cells were pelleted, washed with sterile MilliQ water, and diluted to OD_600_ = 1.85 in sporulation (SPO) media (0.3% potassium acetate, 0.02% raffinose, 40 mg/L adenine, 40 mg/L uracil, 10 mg/L histidine, 10 mg/L leucine and 10 mg/L tryptophan, pH 7). Meiotic cultures were grown at 30°C in flasks that were 10 times the culture volume to ensure maximum aeration.

For two biological replicates using strains containing the *pCUP-IME1/pCUP-IME4* system (strain A33366/UB1007), cells were incubated in SPO media for 2 hours and then 50 mM of CuSO4 was added to induce the expression of *IME1* and *IME4*, which allowed cells to synchronously enter meiosis. Samples were harvested for RNA extraction at the times indicated in Figure 1A. For two biological replicates using strains containing the *GALp-NDT80* (strain A22678/UB140), cells were incubated in SPO media for 5 hours and then 1 uM b-estradiol was added to induce *NDT80* expression, which allowed cells to synchronously exit pachytene. Again, samples were harvested for RNA extraction at the times indicated in Figure 1A.

### RNA extraction

For each meiotic time course, 4 OD_600_ units were pelleted, snap frozen in liquid nitrogen and stored at −80°C. Cells were thawed on ice and then resuspended in 400 µl TES lysis buffer (10 mM Tris pH 7.5, 10 mM EDTA, 0.5% SDS), 400 µL of acid phenol:chloroform:isoamyl alcohol (125:24:1; pH 4.7; Ambion) and 200 uL of acid-washed glass beads (0.5 mm; Sigma). Samples were incubated in a Thermomixer C (Eppendorf) for 30 min at 65°C shaking at 1400 RPM then spun at max speed for 10 min at 4°C. All samples were maintained on ice as the top aqueous phase was transferred to a second tube containing 300 µL of chloroform. Tubes were vortexed for 30 s and the aqueous layer was isolated by spinning at max speed for 5 min at room temperature. The aqueous phase was again moved to a new tube and RNA was then precipitated in 450 µL of isopropanol, 50 µL 3M sodium acetate (NaOAc) pH 5.2. Samples were incubated at −20°C overnight, pelleted, washed with 500 uL of 80% ethanol (diluted with DEPC water), and dried pellets were resuspended in 50 µL DEPC water by incubating in the Thermomixer C for 15 min at 37°C shaking at 1400 RPM.

### Spindle staining and staging

Indirect immunofluorescence was used to stage synchronized meiotic samples as described previously (Kilmartin and Adams, 1984). Spindle morphology was visualized using a rat anti-tubulin antibody (Serotec, Kidlington, UK, diluted 1:100), and anti-rat FITC antibodies (Jackson ImmunoResearch Laboratories, Inc. West Grove, PA, diluted 1:100–200). Spindle number and position relative to DAPI masses were used to differentiate between cells undergoing the first and second meiotic division. Cells were considered in “Metaphase I” when short, bipolar spindles were seen to span a single DAPI mass, “Anaphase I” when elongated spindles were seen to span two DAPI masses, “Metaphase II” when two short, bipolar spindles were detected at each of the two DAPI masses, and “Anaphase II” when elongated spindles were detected at each of the two DAPI masses. Cells that did not fall into any of these categories were counted as “Other”.

### mRNA extraction

Total RNA concentrations were measured on a Qubit 3 (ThermoFisher Scientific) using a Qubit RNA BR Assay Kit (Q10211, ThermoFisher Scientific). Total RNA from timepoints representing starvation, early meiosis, meiosis I, meiosis II, or late meiosis were combined at equal concentrations. mRNA was isolated from each RNA pool using Dynabeads^TM^ Oligo(dT)_25_ (Ambion 61005) according to the provided manual. Briefly, 75µg of total RNA was diluted in 100 µL of distilled DEPC water and 100 µL of Binding Buffer containing 20 mM Tris-HCl pH 7.5, 1.0 M LiCl, and 2 mM EDTA. Each sample was heated to 65°C for 2 minutes, then cooled on ice. 200 µL of Dynabeads^TM^ were washed with an equal volume of Binding Buffer, resuspended in 100 µL of Binding Buffer, and added to each sample. Beads and RNA were incubated on a rotator for 5 min to allow for binding. Beads bound to mRNA were then pelleted using a magnet, the supernatant was removed, and then the samples were resuspended in 200 µL TE (10 mM Tris-HCl pH 7.5 and 0.15 mM EDTA). Beads were pelleted again using a magnet and washed twice with 10 mM Tris-HCl pH 7.5. Beads were then resuspended in 10 µL Tris-HCl and heated to 77°C for 2 minutes. Each sample was then quickly placed onto the magnet and the clear supernatant containing extracted mRNA was removed and stored in low-adhesion Eppendorf tubes at −20°C.

### cDNA library plasmid construction

First strand synthesis was performed on the mRNA pools using a modified protocol from Cloneminer II (ThermoFisher) but with the Superscript reverse transcriptase replaced with the TGIRT reverse transcriptase (Mohr et al., 2013) and primers containing the attB2 site were used. cDNA was isolated using phenol/chloroform/isoamyl alcohol extraction followed by ethanol precipitation. Next, second strand synthesis was performed using T4 DNA polymerase (New England Biolabs) to make blunt ends. cDNA was isolated using phenol/chloroform/isoamyl alcohol extraction followed by ethanol precipitation. The attB1 adapter was ligated onto the libraries using T4 DNA ligase (New England Biolabs). cDNA containing both the attB1 and attB2 sites were isolated using column purification and inserted into pDONR222 using the standard Gateway BP reaction (Katzen, 2007). The cDNA libraries in entry vectors were transformed into ElectroMAX DH10B T1 cells (ThermoFisher) using electroporation, plated, and isolated using a Qiagen Megaprep. cDNA libraries were moved to pUB914 using the standard Gateway LR reaction and cDNA libraries in the pUB914 vector were transformed and isolated in the same way as the cDNA entry vector libraries.

### Yeast transformations with electroporation

Yeast cells were grown in 10 mL of liquid YPD (1% yeast extract, 2% peptone, 2% glucose, 22.4 mg/L uracil, and 80 mg/L tryptophan) overnight shaking at 30°C and then diluted to OD_600_ = 0.2 in 100 mL of YPD. Cultures were grown at 30°C for 3-4 hours until OD_600_ > 0.6. Cells were then spun down in 2 x 50 mL conical tubes at 1,900 g for 2 min at 4°C. Cells were washed twice with 4°C sterile MilliQ water, resuspended in 2 mL of 4°C 1M sorbitol and transferred into 2 x 2 mL Eppendorf tubes. Next, the two tubes of cells were spun at 4°C at 3,200g for 3 min and resuspended in 2 mL of 4°C fresh LiTE solution (1X TE (10 mM Tris-HCl pH 7.9, 1 mM EDTA pH8), 0.1M lithium acetate, 25 mM DTT). Cells were then washed with 2 mL of 4°C 1M sorbitol and resuspended in 250 µL of 4°C 1M sorbitol. Five micrograms of the desired cDNA plasmid library were added to each tube and briefly vortexed and kept on ice. Electroporation was performed on a Bio-Rad Gene Pulser II electroporator with a Capacitance Extender Plus and Pulse Controller Plus system. Fifty microliters of cells were added to a chilled Gene Pulser cuvette with 0.2 cm electrode gap (Bio-Rad) and subjected to a pulse of 1.5 kV, 25 µF, and 100 Ω (pulse time was ∼2.5 ms). Cells were then washed from cuvette using 3 x 1 mL aliquots of YPD and transferred into a 50 mL flask. This process was repeated with all cells and YPD was added to the combined culture up to 50 mL. Electroporated cells were recovered shaking at 30°C for 2 hours, spun down, and resuspended into 500 mL YPD + 320 µg/mL G-418 sulfate (Gibco). After 2 days of growth, OD_600_ was measured, and supernatant was removed such that the cell concentration was increased to 50 OD_600_ units per mL. Five hundred microliters aliquots of cells were then mixed with 500 µL of 30% glycerol in a Cryo-Safe tubes. Aliquots of the cDNA library were stored long term at −80°C.

### Yeast plasmid purification

All yeast samples were frozen in a screw-cap tube with 15% glycerol at −80°C for at least 1 day because a freeze-thaw cycle increased plasmid recovery likely by increasing cell lysis. Frozen cells were thawed at room temperature, pelleted, and resuspended in 250 µL of Qiagen buffer P1 and 200 µL of acid-washed glass beads (0.5 mm; Sigma). Samples were vortexed on high for 5 minutes, 250 µL of Qiagen P2 buffer was added, and tubes were inverted 5 times. Next, 350 µL of Qiagen Q3 buffer was added within 2-3 min of adding P2 to prevent sample degradation. Samples were spun at max speed for 10 minutes and plasmids were purified using a QIAprep spin column according to the Qiagen Miniprep Kit manual. Plasmids were eluted by incubating columns with 50 µL of Qiagen EB buffer for 1 min before spinning on max for 1 min.

### Deep sequencing

For mRNA-seq libraries, 100 ng of polyA-selected mRNA from each meiotic stage was prepared using the NEXTFLEX^TM^ Rapid Directional RNA-seq Kit (NOVA-5138-08; PerkinElmer) according to the provided manual. For DNA-seq libraries, the cDNAs in each plasmid pool were amplified with the KAPA HiFi PCR Kit (KK2101) using SK and KS primers. PCR products were purified using the QIAquick PCR Purification Kit (Qiagen) and the NEXTFLEX^TM^ Rapid DNA-seq Kit (NOVA-5114-03 and NOVA-5144-04; PerkinElmer) was used to prepare 10-50 ng of cDNA. For both mRNA-seq and DNA-seq, all measurements of RNA and DNA were performed on a Qubit 3 (ThermoFisher Scientific) using the appropriate assay kit. AmPure XP beads (A63881, Beckman Coulter) were used for size selection (200-500 bp) and libraries were quantified and quality checked using high sensitivity D1000 ScreenTapes on the Agilent 4200 TapeStation (Agilent Technologies, Inc.). All samples were sequenced by the Vincent J. Coates Genomics Sequencing Laboratory at the University of California, Berkeley using 150 bp single end sequencing on an Illumina Hiseq 4000 or 100 bp single end sequencing on an Illumina Novaseq 6000.

### Sequencing analysis

Sequencing reads were aligned to the SK1 reference genome (from the *Saccharomyces* Genome Resequencing Project (Sanger Institute)) using HISAT2 and transcripts per million reads (TPM) was calculated using STRINGTIE (Pertea et al., 2016).

### Data visualization

For the heatmap in Figure 1B, hierarchical clustering was performed using Cluster 3.0 using uncentered correlation clustering with the centered setting (de Hoon et al., 2004). Data normalization and visualization of the results were performed using Java Treeview (Saldanha, 2004). All plots were made using either Python, Prism 7 (v7.0e) or Microsoft Excel (version 16.48).

### Venn Diagrams

Venn diagrams showing the overlap of 4 or more datasets were created using the InteractiVenn online tool (Heberle et al., 2015). Venn diagrams showing the proportional overlap of 3 datasets were created using the BioVenn online tool (Hulsen et al., 2008).

### Gene set enrichment analysis (GSEA)

Gene Set Enrichment Analysis (GSEA) v4.1.0 [build: 27] was used to compare TPM values for different gene sets across different cDNA libraries (Mootha et al., 2003; Subramanian et al., 2005) The “DNA Replication and Recombination” gene set was created from Figure 2 from Brar et al. (2012). All genes listed in the DNA replication cluster and recombination cluster were combined and any genes that were not present in the TPM tables from this study were removed. The “Ndt80 Cluster” gene set was created by identifying all genes that showed an Ndt80-like gene expression pattern during meiosis using hierarchical clustering (see methods in “Data visualization”) of mRNA data in Cheng and Otto et al. (2018). GSEA was performed on the desktop app with default settings except “Collapse/Remap to gene symbols” was set to “No_Collapse”, “Permutation type” was set to “gene_set”, and the “Max size: exclude larger sets” was set to 800.

### Read density analysis

Gene sizes were calculated from the SK1 reference genome obtained from the *Saccharomyces* Genome Resequencing Project (Sanger Institute). Genes were ordered by size and divided into equal quartiles defined as small, medium, large and extra-large genes. Hybrid reads in the Fastq file (reads that contain cDNA sequence and plasmid backbone sequence) were pulled, the plasmid sequence was trimmed, and the cDNA sequence was added back to the Fastq file to prevent truncation artifacts. Reads were mapped to the SK1 reference genome using HISAT2. The SK1 annotation file was modified to divide each gene into 10 segments and This new annotation file was used to calculate TPM per segment using STRINGTIE (Pertea et al., 2016). TPM was normalized by dividing all segments by the maximum TPM for a given gene and average TPM across different size quartiles was plotted. All analysis and plotting were performed using Python.

### Pipeline for 5’- and 3’- truncation candidates

For all truncation analysis, genes must have an average TPM across the 10 segments greater than 10. Extra-large genes that had extreme 3’-bias in read density in Supplemental Figure 3 were identified with the following criteria: the top 3 largest genes that had a very low TPM across segments 1 and 2, the maximum TPM for segment 10, and segment 10 had a TPM greater than 20. TPM was normalized by dividing all segments by the maximum TPM for a given gene. For 5’ truncation candidates, average normalized TPM of segments 9 and 10 minus the average normalized TPM of segments 1 and had to be greater than 0.33. For 3’ truncations candidates, average normalized TPM of segment 1 and 2 minus average normalized TPM of segments 9 and 10 had to be greater than 0.33. Visual confirmation of gene truncations and read density plots were made using the Integrative Genomics Viewer from the Broad Institute (Robinson et al., 2011).

### Competitive screens

A fresh aliquot of the desired yeast cDNA library was thawed and used to inoculate a YPR (1% yeast extract, 2% peptone, 2% raffinose, 22.4 mg/L uracil, and 80 mg/L tryptophan) + 320 μg G418 culture at OD_600_ = 0.2. The yeast library was recovered at 30°C for 8 hours and then 2 million cells were washed with YPGR media (1% yeast extract, 2% peptone, 2% raffinose, 2% galactose, 22.4 mg/L uracil, and 80 mg/L tryptophan) and 2 million cells were washed with YPD media (1% yeast extract, 2% peptone, 2% glucose, 22.4 mg/L uracil, and 80 mg/L tryptophan). Cells were then used to inoculate a 50 mL culture with inducing YPGR + 320 μg G418 media or uninducing YPD + 320 μg G418 media. Cells were grown at 30°C for 24 hours and 25 OD_600_ units were spun down and resuspended in 15% glycerol and stored at −80°C until processed for sequencing. Saturated cells were re-diluted 1 in 1000 in the appropriate media and this cycle was continued for 5 days with samples harvested every 24 h.

### Monotonicity z-scores

To identify cDNAs that become enriched or disenriched within a given library, we adapted a permutation-based method to measure the monotonic change in TPM for each gene over time (Taliaferro et al., 2016). For each gene, TPM values were ordered chronologically and pair-wise comparisons between a given timepoint and each successive timepoint were performed using a T-test. A tally of the number of comparisons representing a significant (p-value < 0.05) increase (i) or decrease (d) across three replicates were calculated, and these tallies were used to calculate δ using the formula δ = i - d. To test for statistical significance, the timepoints for each gene were scrambled and a new δ value was calculated. This process was repeated 1000 times to generate a null distribution of δ values, which was used to calculate a mean (μ) and standard deviation (σ) value. A monotonicity z-score (MZ-score) for each gene was then calculated using the formula MZ = (δ – μ)/σ.

### Plasmid construction

A KAPA HiFi PCR Kit (KK2101) was used to amplify each gene from genomic DNA isolated from wild-type SK1 (UB13) using primers that would add attB1 to the 5’ end of the gene and attB2 to the 3’ end of the gene (Supplementary Table 9). PCR products were purified using a QIAquick PCR Purification Kit (Qiagen) and each gene was inserted into an entry vector (pDONR222) using the standard BP reaction outlined in the Gateway Cloning Protocol. Each entry vector was amplified in DH5alpha cells and isolated with a Qiagen Miniprep Kit. Insertion was confirmed by running 10 ng of plasmid onto an 1% agarose gel. Each gene was then moved to the galactose-inducible destination plasmid (pUB914) using the standard LR reaction outlined in the Gateway Cloning Protocol. Each destination vector was amplified in DH5alpha cells, purified with the Qiagen Miniprep Kit and the entire plasmid was sequenced at Octant using their Octopus platform (https://www.octant.bio/blog/2019/9/29/octopus).

### Gene ontology enrichment analysis

All gene ontology (GO) enrichment was performed using g:profiler (Raudvere et al., 2019) or Yeast Mine (https://yeastmine.yeastgenome.org/) using default settings except terms that were greater than 1000 genes were omitted.

### Competitive growth assay

Each plasmid was transformed into the appropriate strain using standard yeast techniques. Three individual colonies representing three biological replicates were used to inoculate 1 mL of YPR + 320 μg/mL G418 grown in a 2 mL well with a single glass bead in a 96-well culture box. Culture boxes were grown shaking overnight at 30°C for at least 16 h to ensure saturation of growth. Ten microliters of saturated culture from each of the competing strains were transferred to 1 mL YPGR + 320 μg/mL G418 to induce gene expression. Culture boxes were grown for another 24 h shaking at 30°C and 10 million cells were fixed for flow cytometry. Saturated cells in the culture box were then re-diluted 1 in 100 in 1 mL YPGR + 320 μg/mL G418 for a second day of growth and a “Day 2” sample was fixed for flow cytometry.

### Formaldehyde fixation

Cells expressing *PGK1::eGFP* were fixed by adding the appropriate volume of 37% formaldehyde directly into the media (final concentration was 3.7%). Tubes were inverted 5-6 times and incubated at room temperature for 15 min. Cells were then washed in 100 mM potassium phosphate, pH 6.4, and stored at 4°C in KPi Sorbitol soluton (100 mM potassium phosphate, pH 7.5, 1.2 M sorbitol). Fluorescence measurements using a flow cytometer were performed within 3 days of formaldehyde fixation.

### Flow cytometry

Formaldehyde fixed cells stored in KPi sorbitol buffer were spun down and resuspended in 1X PBS pH 7 to a final density of 1 OD_600_ unit per mL. Cells were passed through a cell-strainer cap on a 5 mL polystyrene round-bottom tube (FALCON 352235) and GFP signal was measured on a BD LSR Fortessa (BD Biosciences) at the Flow Cytometry Facility at the University of California, Berkeley.

### Non-competitive growth curves

Each plasmid was transformed into UB20279 using standard yeast techniques. Three individual colonies representing three biological replicates were used to inoculate 1 mL of YPR + 320 μg/mL G418 grown in a 2 mL well with a single glass bead in a 96-well culture box. Culture boxes were grown shaking overnight at 30°C for at least 16 h to ensure saturation of growth. Fifty microliters of saturated culture was mixed with 950 μL of YPGR or YPD and 200 μL was transferred to a 96-well flat bottom plate. A single OD_600_ measurement was taken on a TECAN Spark microplate reader and used to calculate the volume required to dilute cultures to 200 μL of OD_600_ = 0.05. The TECAN Spark microplate reader was then used to measure OD_600_ every 15 min for 24 – 48 hr at 30°C with shaking in between measurements. Spark Control Magellan^TM^ software (version 2.3) was used for data acquisition.

### Data availability

Sequencing data produced from this study can be found at the NCBI Gene Expression Omnibus (GEO) with the accession number TBD. All code used for identifying 5’- and 3’- truncation candidates as well as for calculating MZ-scores for the competitive screens are available at: https://github.com/sudmantlab/SingYeastcDNAScreening.

## ACKNOWLEDGEMENTS

We thank Grant King and Jay Goodman for their comments on this manuscript as well as all members of the Ünal, Brar, and Sudmant labs for scientific discussions for this project. We also thank Leon Chan and Gabrielle Bostwick from Octant for facilitating plasmid sequencing using the Octopus platform. This research was supported by funds from the Pew Charitable Trusts (00027344), Damon Runyon Cancer Research Foundation (35-15), National Institutes of Health (DP2 AG055946-01) and Curci Foundation to E.Ü and the National Institutes of Health (R35 GM134886) to G.A.B. T.L.S. was funded in part by a Postdoctoral Fellowship (PDF-532943-2019) from the Natural Sciences and Engineering Research Council of Canada (NSERC) and I.H. was funded by an Erwin Schrödinger Fellowship.

## SUPPLEMENTAL FIGURE LEGENDS

**Supplemental Figure 1.**
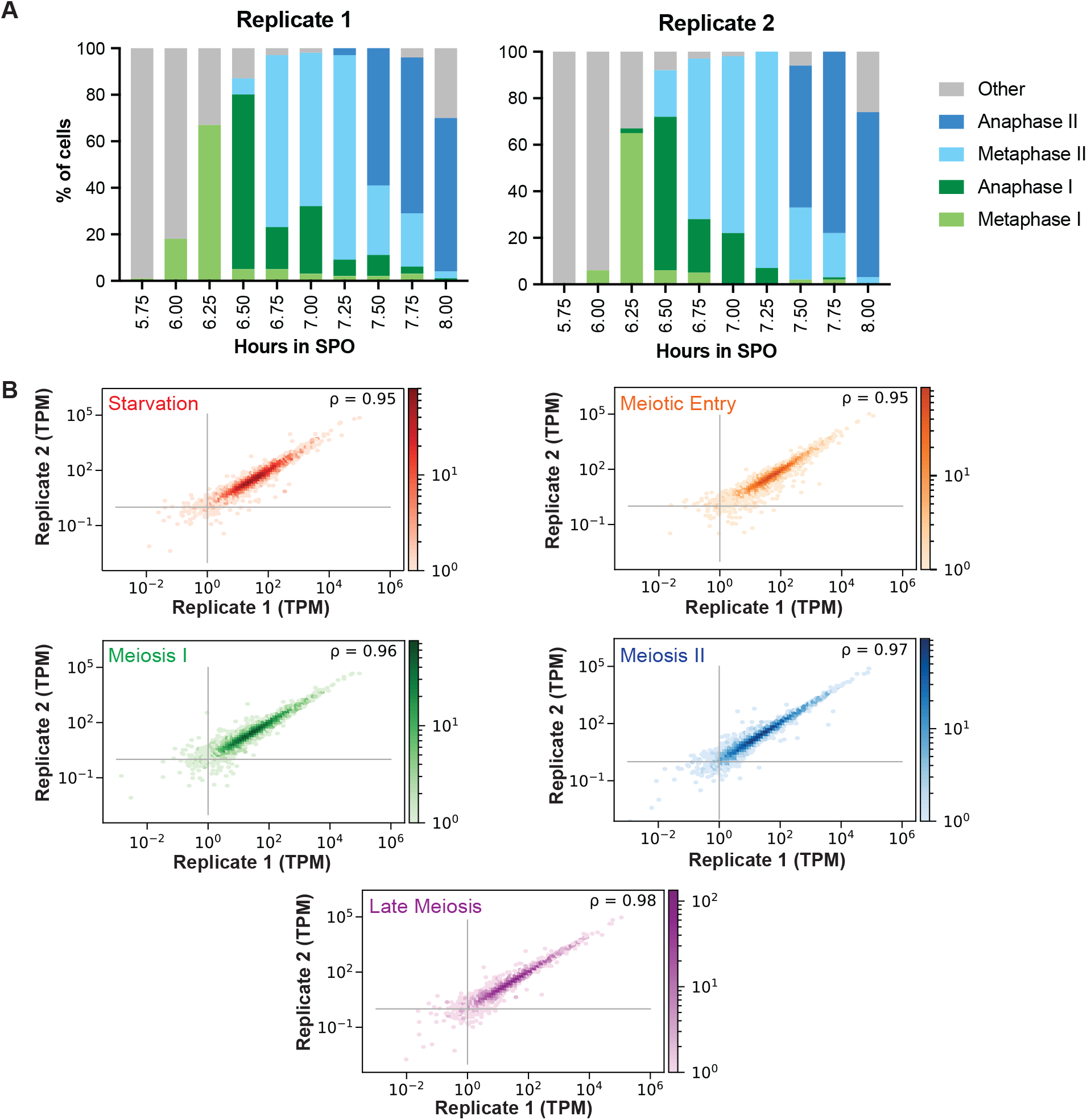
Replicates of stage-specific meiotic RNA pools are highly reproducible. **(A)** Cells expressing a galactose-inducible *NDT80* allele were grown in sporulation media (SPO) for 5 hours to arrest cells in pachytene. *NDT80* expression was then induced with β-estradiol and samples were taken at the indicated time points. Spindles were then stained to quantify meiotic progression in each replicate. **(B)** The mRNA from the indicated pools were sequenced and replicates were plotted for comparison. The spearman’s rank order correlation coefficient is displayed at the top right corner of each graph. The plot density is displayed on the right. TPM, transcripts per million reads.

**Supplemental Figure 2.**
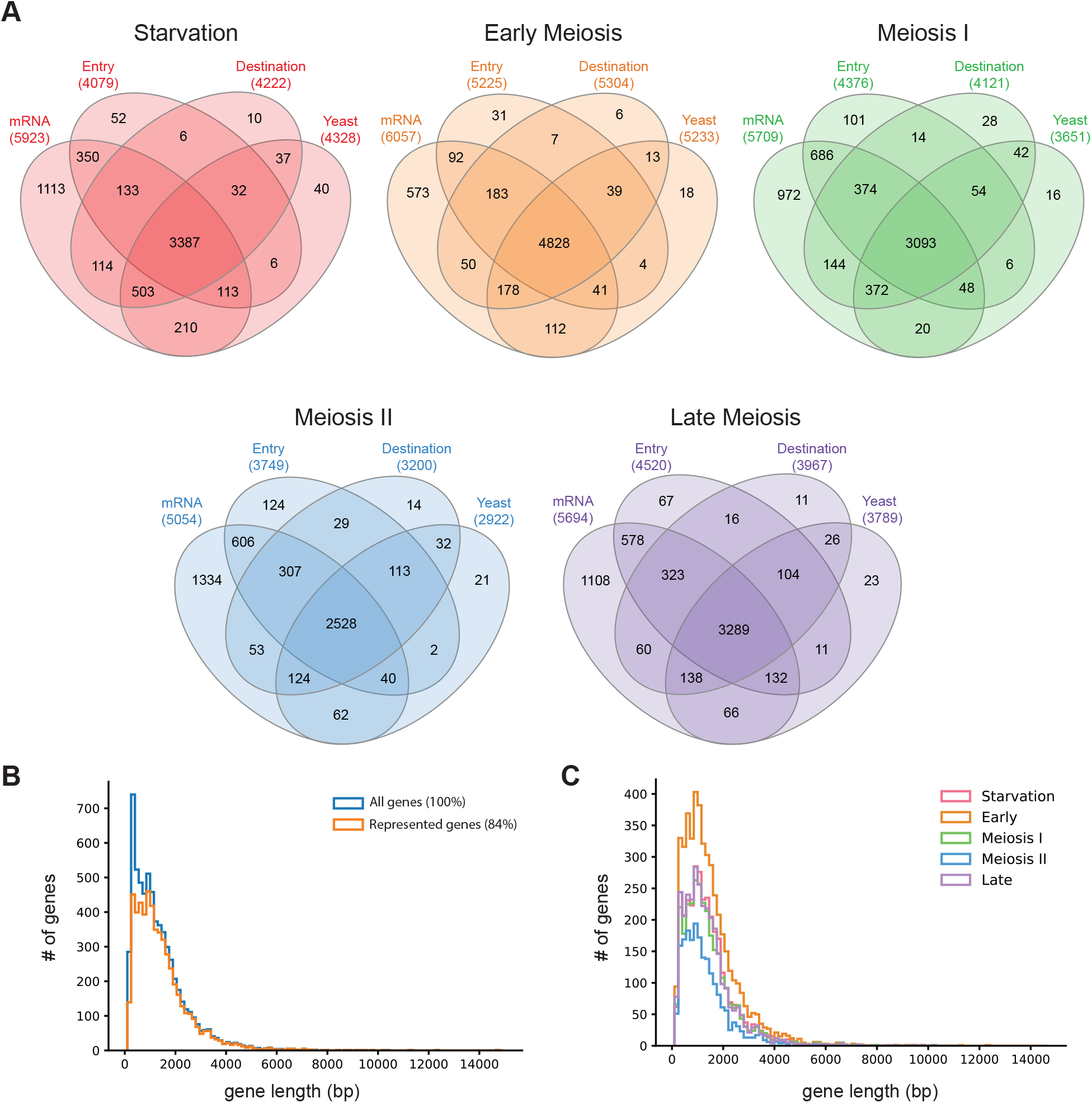
Estimating the number of genes represented in each meiotic library. **(A)** Genes that have a TPM value greater than 5 in the mRNA, entry vector, destination vector, and yeast pools were overlapped in Venn Diagrams for the indicated libraries. **(B)** Comparison of gene length in base pairs (bp) across all genes in the yeast genome (100%) and genes represented in at least one of the five meiotic cDNA libraries (84%) across 150 bp bins. **(C)** The distribution of gene length in base pairs (bp) between each meiotic cDNA library across 150 bp bins.

**Supplemental Figure 3.**
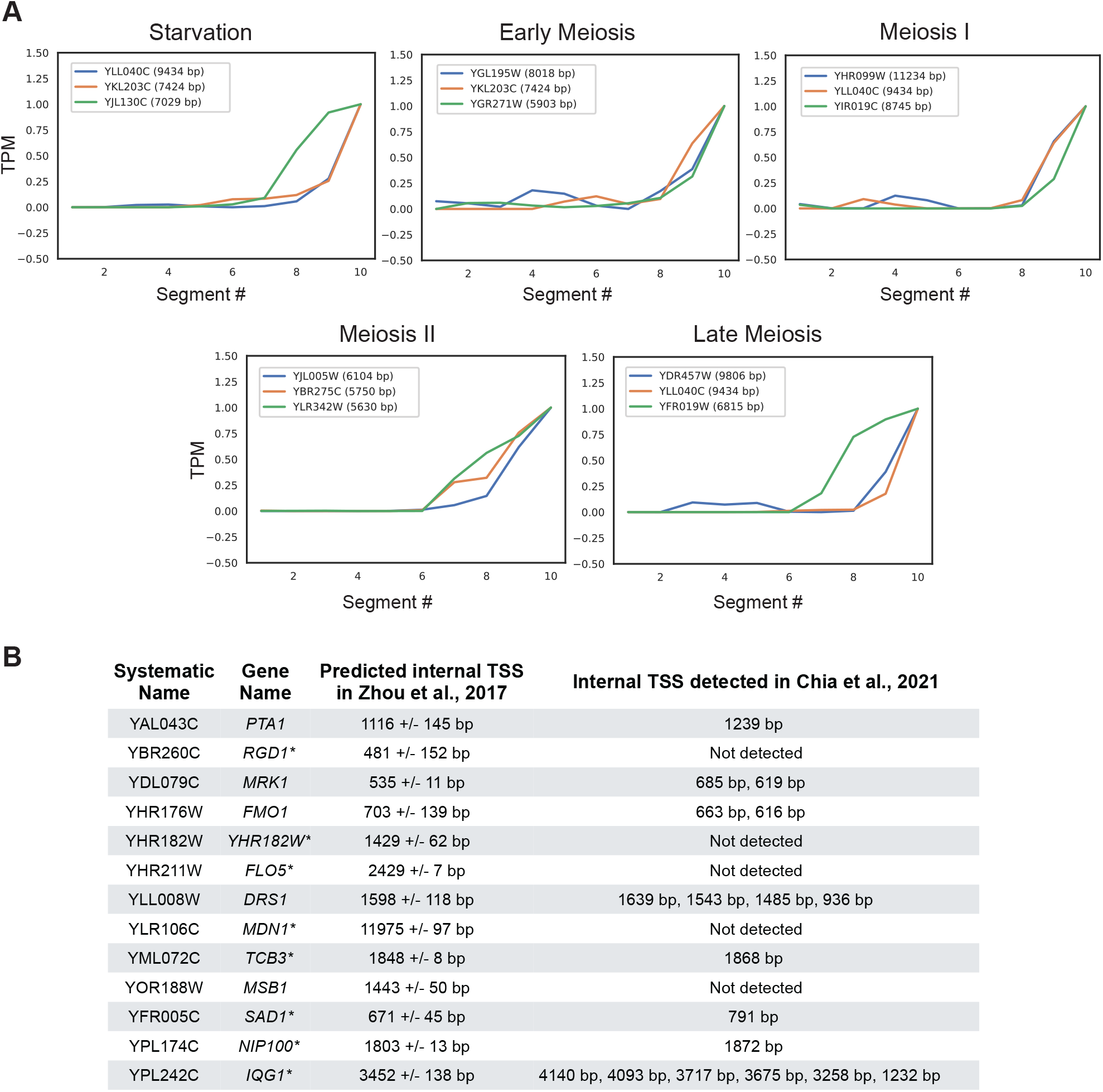
Reverse transcription may introduce 3’-bias in extra-large genes. **(A)** The three largest genes from each cDNA library that had a very low TPM for segments 1-3 (average TPM for segments 1-3 less than 1.5) and the maximum TPM for segment 10 (TPM greater than 20). The gene name and length in base pairs (bp) is indicated in the legend for each graph. **(B)** A list of 13 intragenic transcript isoforms found in this study that were previously identified computationally in Zhou et al. (2017). All 5’-truncations were confirmed by visual inspection of read density plots. Genes marked with an asterisk (*) were confirmed by visual inspection but did not reach the threshold for our computational analysis. The distance of the intragenic transcription start site (TSSs) from the canonical TSS according to Zhou et al. (2017) and Chia et al. (2021) are also shown.

**Supplemental Figure 4.**
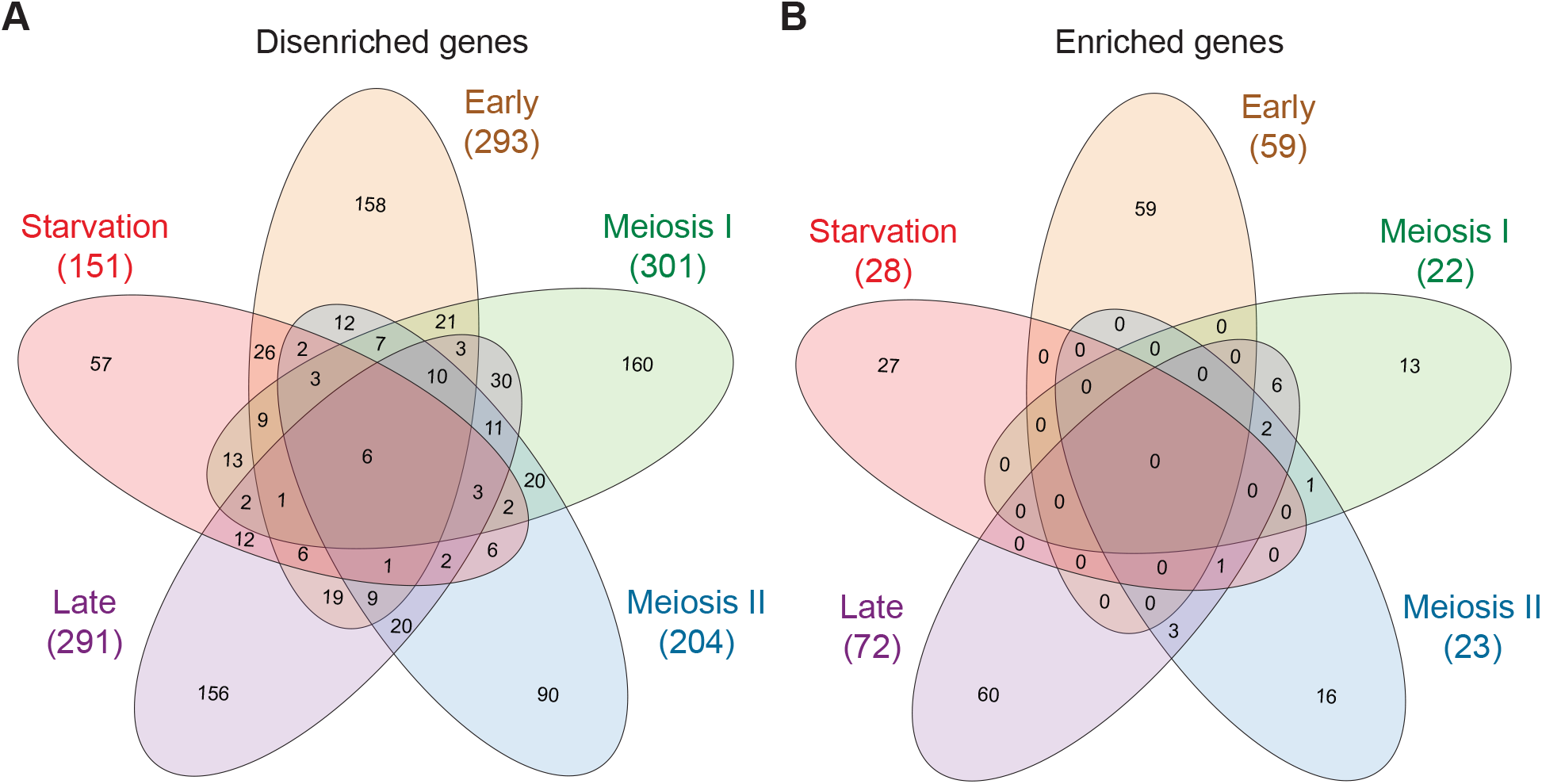
Number of genes that are beneficial or detrimental to competitive growth in each cDNA library. **(A)** Venn diagram showing the overlap of cDNAs that disenrich in the population during the competitive screening assay (MZ-score < −1.8) in each of the libraries. **(B)** Venn diagram showing the overlap of cDNAs that enrich in the population during the competitive screening assay (MZ-score > 1.8) in each of the libraries.

## REFERENCES

Almagro Armenteros, J.J., Salvatore, M., Emanuelsson, O., Winther, O., von Heijne, G., Elofsson, A., and Nielsen, H. (2019). Detecting sequence signals in targeting peptides using deep learning. Life Sci. Alliance 2, e201900429.

Arita, Y., Kim, G., Li, Z., Friesen, H., Turco, G., Wang, R.Y., Climie, D., Usaj, M., Hotz, M., Stoops, E., et al. (2021). A genome-scale yeast library with inducible expression of individual genes. BioRxiv 2020.12.30.424776.

Baillie, K., Hoon, M.J.L. De, Lassmann, T., Itoh, M., Summers, K.M., Suzuki, H., and Daub, C.O. (2014). A promoter-level mammalian expression atlas. Nature 507, 462– 470.

Bannai, H., Tamada, Y., Maruyama, O., Nakai, K., and Miyano, S. (2002). Extensive feature detection of N-terminal protein sorting signals. Bioinformatics 18, 298–305.

Ben-Ari, G., Zenvirth, D., Sherman, A., David, L., Klutstein, M., Lavi, U., Hillel, J., and Simchen, G. (2006). Four Linked Genes Participate in Controlling Sporulation Efficiency in Budding Yeast. PLoS Genet. 2, e195.

Benjamin, K.R., Zhang, C., Shokat, K.M., and Herskowitz, I. (2003). Control of landmark events in meiosis by the CDK Cdc28 and the meiosis-specific kinase Ime2. Genes Dev. 17, 1524–1539.

Berchowitz, L.E., Gajadhar, A.S., van Werven, F.J., De Rosa, A.A., Samoylova, M.L., Brar, G.A., Xu, Y., Xiao, C., Futcher, B., Weissman, J.S., et al. (2013). A developmentally regulated translational control pathway establishes the meiotic chromosome segregation pattern. Genes Dev. 27, 2147–2163.

Bolcun-Filas, E., and Handel, M.A. (2018). Meiosis: the chromosomal foundation of reproduction. Biol. Reprod. 99, 112–126.

Boos, F., Labbadia, J., and Herrmann, J.M. (2020). How the Mitoprotein-Induced Stress Response Safeguards the Cytosol: A Unified View. Trends Cell Biol. 30, 241–254.

Brar, G.A., Yassour, M., Friedman, N., Regev, A., Ingolia, N.T., and Weissman, J.S. (2012). High-resolution view of the yeast meiotic program revealed by ribosome profiling. Science 335, 552–557.

Carlile, T.M., and Amon, A. (2008). Meiosis I is established through division-specific translational control of a cyclin. Cell 133, 280–291.

Chen, J., Tresenrider, A., Chia, M., McSwiggen, D.T., Spedale, G., Jorgensen, V., Liao, H., van Werven, F.J., and Ünal, E. (2017). Kinetochore inactivation by expression of a repressive mRNA. Elife 6, 1–31.

Cheng, Z., Otto, G.M., Powers, E.N., Keskin, A., Mertins, P., Carr, S.A., Jovanovic, M., and Brar, G.A. (2018). Pervasive, Coordinated Protein-Level Changes Driven by Transcript Isoform Switching during Meiosis. Cell 172, 910–914.e16.

Chia, M., Tresenrider, A., Chen, J., Spedale, G., Jorgensen, V., Ünal, E., and van Werven, F.J. (2017). Transcription of a 5’ extended mRNA isoform directs dynamic chromatin changes and interference of a downstream promoter. Elife 6, 1–23.

Chia, M., Li, C., Marques, S., Pelechano, V., Luscombe, N.M., and van Werven, F.J. (2021). High-resolution analysis of cell-state transitions in yeast suggests widespread transcriptional tuning by alternative starts. Genome Biol. 22, 46.

Chu, S., DeRisi, J., Eisen, M., Mulholland, J., Botstein, D., Brown, P.O., and Herskowitz, I. (1998). The transcriptional program of sporulation in budding yeast. Science 282, 699–705.

Claros, M.G., and Vincens, P. (1996). Computational method to predict mitochondrially imported proteins and their targeting sequences. Eur. J. Biochem. 241, 779–786.

Demircioğlu, D., Cukuroglu, E., Kindermans, M., Nandi, T., Calabrese, C., Fonseca, N.A., Kahles, A., Lehmann, K.-V., Stegle, O., Brazma, A., et al. (2019). A Pan-cancer Transcriptome Analysis Reveals Pervasive Regulation through Alternative Promoters. Cell 178, 1465–1477.e17.

Douglas, A.C., Smith, A.M., Sharifpoor, S., Yan, Z., Durbic, T., Heisler, L.E., Lee, A.Y., Ryan, O., Göttert, H., Surendra, A., et al. (2012). Functional Analysis With a Barcoder Yeast Gene Overexpression System. G3 Genes|Genomes|Genetics 2, 1279–1289.

Eastwood, M.D., and Meneghini, M.D. (2015). Developmental Coordination of Gamete Differentiation with Programmed Cell Death in Sporulating Yeast. Eukaryot. Cell 14, 858–867.

Eastwood, M.D., Cheung, S.W.T., and Meneghini, M.D. (2013). Programmed nuclear destruction in yeast Self-eating by vacuolar lysis. Autophagy 9, 263–265.

Eisenberg, A.R., Higdon, A.L., Hollerer, I., Fields, A.P., Jungreis, I., Diamond, P.D., Kellis, M., Jovanovic, M., and Brar, G.A. (2020). Translation Initiation Site Profiling Reveals Widespread Synthesis of Non-AUG-Initiated Protein Isoforms in Yeast. Cell Syst. 11, 145–160.e5.

Freese, E.B., Chu, M.I., and Freese, E. (1982). Initiation of Yeast sporulation by partial carbon, nitrogen, or phosphate deprivation. J. Bacteriol. 149, 840–851.

Fukasawa, Y., Tsuji, J., Fu, S.-C., Tomii, K., Horton, P., and Imai, K. (2015). MitoFates: Improved Prediction of Mitochondrial Targeting Sequences and Their Cleavage Sites*. Mol. Cell. Proteomics 14, 1113–1126.

Gelperin, D.M. (2005). Biochemical and genetic analysis of the yeast proteome with a movable ORF collection. Genes Dev. 19, 2816–2826.

Goodman, J.S., King, G.A., and Ünal, E. (2020). Cellular quality control during gametogenesis. Exp. Cell Res. 396, 112247.

Heberle, H., Meirelles, G.V., da Silva, F.R., Telles, G.P., and Minghim, R. (2015). InteractiVenn: a web-based tool for the analysis of sets through Venn diagrams. BMC Bioinformatics 16, 169.

Ho, C.H., Magtanong, L., Barker, S.L., Gresham, D., Nishimura, S., Natarajan, P., Koh, J.L.Y., Porter, J., Gray, C.A., Andersen, R.J., et al. (2009). A molecular barcoded yeast ORF library enables mode-of-action analysis of bioactive compounds. Nat. Biotechnol. 27, 369–377.

Hollerer, I., Curk, T., Haase, B., Benes, V., Hauer, C., Neu-Yilik, G., Bhuvanagiri, M., Hentze, M.W., and Kulozik, A.E. (2016). The differential expression of alternatively polyadenylated transcripts is a common stress-induced response mechanism that modulates mammalian mRNA expression in a quantitative and qualitative fashion. RNA 22, 1441–1453.

de Hoon, M.J.L., Imoto, S., Nolan, J., and Miyano, S. (2004). Open source clustering software. Bioinformatics 20, 1453–1454.

Hu, Y., Rolfs, A., Bhullar, B., Murthy, T.V.S., Zhu, C., Berger, M.F., Camargo, A.A., Kelley, F., McCarron, S., Jepson, D., et al. (2007). Approaching a complete repository of sequence-verified protein-encoding clones for Saccharomyces cerevisiae. Genome Res. 17, 536–543.

Hulsen, T., de Vlieg, J., and Alkema, W. (2008). BioVenn – a web application for the comparison and visualization of biological lists using area-proportional Venn diagrams. BMC Genomics 9, 488.

Kassir, Y., Granot, D., and Simchen, G. (1988). IME1, a positive regulator gene of meiosis in S. cerevisiae. Cell 52, 853–862.

Katzen, F. (2007). Gateway ® recombinational cloning: a biological operating system. Expert Opin. Drug Discov. 2, 571–589.

Kilmartin, J. V., and Adams, A.E.M. (1984). Structural rearrangements of tubulin and actin during the cell cycle of the yeast Saccharomyces. J. Cell Biol. 98, 922–933.

King, G.A., and Ünal, E. (2020). The dynamic nuclear periphery as a facilitator of gamete health and rejuvenation. Curr. Genet. 66, 487–493.

King, G.A., Goodman, J.S., Schick, J.G., Chetlapalli, K., Jorgens, D.M., McDonald, K.L., and Ünal, E. (2019). Meiotic cellular rejuvenation is coupled to nuclear remodeling in budding yeast. Elife 8, 1–32.

Lardenois, A., Liu, Y., Walther, T., Chalmel, F., Evrard, B., Granovskaia, M., Chu, A., Davis, R.W., Steinmetz, L.M., and Primig, M. (2011). Execution of the meiotic noncoding RNA expression program and the onset of gametogenesis in yeast require the conserved exosome subunit Rrp6. Proc. Natl. Acad. Sci. 108, 1058–1063.

Marston, A.L., and Amon, A. (2004). Meiosis: cell-cycle controls shuffle and deal. Nat. Rev. Mol. Cell Biol. 5, 983–997.

Merlini, L., Dudin, O., and Martin, S.G. (2013). Mate and fuse: how yeast cells do it. Open Biol. 3, 130008.

Mohr, S., Ghanem, E., Smith, W., Sheeter, D., Qin, Y., King, O., Polioudakis, D., Iyer, V.R., Hunicke-Smith, S., Swamy, S., et al. (2013). Thermostable group II intron reverse transcriptase fusion proteins and their use in cDNA synthesis and next-generation RNA sequencing. RNA 19, 958–970.

Mootha, V.K., Lindgren, C.M., Eriksson, K.-F., Subramanian, A., Sihag, S., Lehar, J., Puigserver, P., Carlsson, E., Ridderstråle, M., Laurila, E., et al. (2003). PGC-1α-responsive genes involved in oxidative phosphorylation are coordinately downregulated in human diabetes. Nat. Genet. 34, 267–273.

Neiman, A.M. (2005). Ascospore Formation in the Yeast Saccharomyces cerevisiae. Microbiol. Mol. Biol. Rev. 69, 565–584.

Neiman, A.M. (2011). Sporulation in the budding yeast Saccharomyces cerevisiae. Genetics 189, 737–765.

Otto, G.M., Cheunkarndee, T., Leslie, J.M., and Brar, G.A. (2021). Programmed ER fragmentation drives selective ER inheritance and degradation in budding yeast meiosis. BioRxiv 2021.02.12.430990.

Pelechano, V., Wei, W., and Steinmetz, L.M. (2013). Extensive transcriptional heterogeneity revealed by isoform profiling. Nature 497, 127–131.

Pertea, M., Kim, D., Pertea, G.M., Leek, J.T., and Salzberg, S.L. (2016). Transcript-level expression analysis of RNA-seq experiments with HISAT, StringTie and Ballgown. Nat. Protoc. 11, 1650–1667.

Raudvere, U., Kolberg, L., Kuzmin, I., Arak, T., Adler, P., Peterson, H., and Vilo, J. (2019). g:Profiler: a web server for functional enrichment analysis and conversions of gene lists (2019 update). Nucleic Acids Res. 47, W191–W198.

Reyes, A., and Huber, W. (2018). Alternative start and termination sites of transcription drive most transcript isoform differences across human tissues. Nucleic Acids Res. 46, 582–592.

Robinson, J.T., Thorvaldsdóttir, H., Winckler, W., Guttman, M., Lander, E.S., Getz, G., and Mesirov, J.P. (2011). Integrative genomics viewer. Nat. Biotechnol. 29, 24–26.

Rousseau, P., Halvorson, H.O., Bulla, L.A., and Julian, G. St. (1972). Germination and Outgrowth of Single Spores of Saccharomyces cerevisiae Viewed by Scanning Electron and Phase-Contrast Microscopy. J. Bacteriol. 109, 1232–1238.

Saldanha, A.J. (2004). Java Treeview--extensible visualization of microarray data. Bioinformatics 20, 3246–3248.

Sawyer, E.M., Joshi, P.R., Jorgensen, V., Yunus, J., Berchowitz, L.E., and Ünal, E. (2019). Developmental regulation of an organelle tether coordinates mitochondrial remodeling in meiosis. J. Cell Biol. 218, 559–579.

Sopko, R., Huang, D., Preston, N., Chua, G., Papp, B., Kafadar, K., Snyder, M., Oliver, S.G., Cyert, M., Hughes, T.R., et al. (2006). Mapping pathways and phenotypes by systematic gene overexpression. Mol. Cell 21, 319–330.

Subramanian, A., Tamayo, P., Mootha, V.K., Mukherjee, S., Ebert, B.L., Gillette, M.A., Paulovich, A., Pomeroy, S.L., Golub, T.R., Lander, E.S., et al. (2005). Gene set enrichment analysis: a knowledge-based approach for interpreting genome-wide expression profiles. Proc. Natl. Acad. Sci. U. S. A. 102, 15545–15550.

Suda, Y., Nakanishi, H., Mathieson, E.M., and Neiman, A.M. (2007). Alternative Modes of Organellar Segregation during Sporulation in Saccharomyces cerevisiae. Eukaryot. Cell 6, 2009–2017.

Taliaferro, J.M., Lambert, N.J., Sudmant, P.H., Dominguez, D., Merkin, J.J., Alexis, M.S., Bazile, C., and Burge, C.B. (2016). RNA Sequence Context Effects Measured In Vitro Predict In Vivo Protein Binding and Regulation. Mol. Cell 64, 294–306.

Tang, H.-L., Yeh, L.-S., Chen, N.-K., Ripmaster, T., Schimmel, P., and Wang, C.-C. (2004). Translation of a Yeast Mitochondrial tRNA Synthetase Initiated at Redundant non-AUG Codons. J. Biol. Chem. 279, 49656–49663.

Tresenrider, A., Morse, K., Jorgensen, V., Chia, M., Liao, H., Jacobus van Werven, F., and Ünal, E. (2021). Integrated genomic analysis reveals key features of long undecoded transcript isoform-based gene repression. Mol. Cell 1–15.

Unal, E., Kinde, B., and Amon, A. (2011). Gametogenesis eliminates age-induced cellular damage and resets life span in yeast. Science 332, 1554–1557.

Xia, Z., Donehower, L.A., Cooper, T.A., Neilson, J.R., Wheeler, D.A., Wagner, E.J., and Li, W. (2014). Dynamic analyses of alternative polyadenylation from RNA-seq reveal a 3’-UTR landscape across seven tumour types. Nat. Commun. 5, 5274.

Xu, L., Ajimura, M., Padmore, R., Klein, C., and Kleckner, N. (1995). NDT80, a meiosis-specific gene required for exit from pachytene in Saccharomyces cerevisiae. Mol. Cell. Biol. 15, 6572–6581.

Yang, S.W., Li, L., Connelly, J.P., Porter, S.N., Kodali, K., Gan, H., Park, J.M., Tacer, K.F., Tillman, H., Peng, J., et al. (2020). A Cancer-Specific Ubiquitin Ligase Drives mRNA Alternative Polyadenylation by Ubiquitinating the mRNA 3 End Processing Complex. Mol. Cell 77, 1206–1221.e7.

Yoshikawa, K., Tanaka, T., Ida, Y., Furusawa, C., Hirasawa, T., and Shimizu, H. (2011). Comprehensive phenotypic analysis of single-gene deletion and overexpression strains of Saccharomyces cerevisiae. Yeast 28, 349–361.

Zhou, S., Sternglanz, R., and Neiman, A.M. (2017). Developmentally regulated internal transcription initiation during meiosis in budding yeast. PLoS One 12, e0188001.

